# Stitching and registering highly multiplexed whole slide images of tissues and tumors using ASHLAR

**DOI:** 10.1101/2021.04.20.440625

**Authors:** Jeremy L. Muhlich, Yu-An Chen, Clarence Yapp, Douglas Russell, Sandro Santagata, Peter K Sorger

**Author notes:** Peter Sorger, Warren Alpert 440, 200 Longwood Avenue, Harvard Medical School, Boston MA 02115, cc.

## Abstract

**Motivation:** Stitching microscope images into a mosaic is an essential step in the analysis and visualization of large biological specimens, particularly human and animal tissues. Recent approaches to highly-multiplexed imaging generate high-plex data from sequential rounds of lower-plex imaging. These multiplexed imaging methods promise to yield precise molecular single-cell data and information on cellular neighborhoods and tissue architecture. However, attaining mosaic images with single-cell accuracy requires robust image stitching and image registration capabilities that are not met by existing methods.

**Results:** We describe the development and testing of ASHLAR, a Python tool for coordinated stitching and registration of 10^3^ or more individual multiplexed images to generate accurate whole-slide mosaics. ASHLAR reads image formats from most commercial microscopes and slide scanners, and we show that it performs better than existing open source and commercial software. ASHLAR outputs standard OME-TIFF images that are ready for analysis by other open-source tools and recently developed image analysis pipelines.

**Availability and implementation:** ASHLAR is written in Python and available under an MIT license at https://github.com/labsyspharm/ashlar. An informational website with user guides and test data is available at https://labsyspharm.github.io/ashlar/.

## INTRODUCTION

Multiple approaches have been described for performing 20-60 plex subcellular resolution microscopy on normal and diseased tissues for research and diagnostic purposes (Angelo et al., 2014; Gerdes et al., 2013; Giesen et al., 2014; Goltsev et al., 2018; Lin et al., 2018; Tsujikawa et al., 2017). These methods make it possible to image differentiation markers, signaling proteins, cell cycle regulators, oncogenes, and drug targets in a preserved tissue context. The resulting data can be processed to determine the molecular and physical relationships of cells within the tissue to each other, to the local vasculature, and to the noncellular components within connective tissue or basement membranes. Research has shown that spatial profiling by highly-multiplexed microscopy can reveal features of normal and diseased tissues and their responses to therapy that cannot be discerned in other ways (Färkkilä et al., 2020; Goltsev et al., 2018; Launonen et al., 2022; Schürch et al., 2020; Wagner et al., 2019). For this reason, multiplexed spatial profiling of proteins and mRNA is the cornerstone of large scale atlasing projects such as the Human Cell Atlas (Regev et al., 2017), NIH HuBMAP consortium (HuBMAP Consortium, 2019), and NCI Human Tumor Atlas Network (HTAN) (Rozenblatt-Rosen et al., 2020). Such atlases promise to fundamentally advance understanding of tissue development and physiology and improve how diseases are diagnosed and individual patients matched to optimal therapies.

Highly multiplexed imaging of proteins in tissues uses antibodies to detect specific antigens, building on 80 years of experience with immunohistochemistry in research and diagnostic settings (Wick, 2012). Methods such as MxIF, CyCIF, CODEX, 4i, and mIHC use conventional fluorescence and brightfield microscopes whereas MIBI and IMC vaporize specimens with ion beams or lasers followed by atomic mass spectrometry (Angelo et al., 2014; Gerdes et al., 2013; Giesen et al., 2014; Goltsev et al., 2018; Gut et al., 2018, 2018; Lin et al., 2018; Tsujikawa et al., 2017). Approaches to imaging nucleic acids are based on hybridization (Chen et al., 2015; Lee et al., 2014) and sequencing (Ståhl et al., 2016). Some methods require frozen samples, but methods that use Formaldehyde Fixed Paraffin Embedded (FFPE) specimens - the sample type universally acquired for diagnostic purposes - can tap into large archives of human biopsy and resection specimens (Burger et al., 2021).

Existing imaging methods differ in resolution, field of view and number of distinct antigens or genes that can be detected (the assay “plex”). Most immunofluorescence methods (e.g. MxIF, CyCIF, mIHC, CODEX and Immuno-SABER) are cyclic approaches in which high-plex data are generated by repeated acquisition of lower-plex images, each of which has 2-6 channels of information. Each channel represents an image acquired with excitation and emission filters matching one antibody or oligo-coupled fluorophore. As such, cyclic imaging makes it possible to optimally exploit the optical properties of fluorescence microscopes while interrogating 60 or more distinct antigens from a single specimen. Conventional research-grade fluorescence microscopes can acquire data from up to 6 different channels, at resolutions down 0.25 µm (laterally) which makes detailed analysis of intracellular structures possible. “Slide scanners” are microscopes equipped with rigid slide holders that move in X and Y and use non-immersion (air) objectives to rapidly move across the specimen. At resolutions sufficient for subcellular imaging, collecting data from a whole slide involves acquiring an array of multiple image “tiles” (10^3^ or more for a large specimen of 6 cm^2^). Thus, each tile is a multi-wavelength megapixel-scale image that represents a different lateral (x, y) stage position. The number of wavelengths in each tile, the number of tiles, and the number of imaging “cycles” (each of which involves acquisition of a full set of tiles), differs with the microscope and the multiplexing technology. However, it is universally true that tiles from all cycles must be merged accurately into a single high-plex “mosaic” image.

High-plex mosaic images represent the key “Level 2 or 3” data type for all subsequent visualization and quantitative data analysis. The data level concept was introduced by dbGAP for genomics (Tryka et al., 2014) and its implementation to tissue imaging is described in detail in the MITI guidelines (Schapiro et al., 2022a). In this context, “data levels” denote different degrees of data processing, with Level 1 corresponding to single, raw image tiles, Levels 2 data to stitched, illumination corrected mosaics and Level 3 to mosaic images that have also been subjected to manual or automated quality control to improve interpretability and accuracy.

It is increasingly clear that the greatest challenges in the acquisition and analysis of high-plex image data lies not in image acquisition per se, but in the subsequent image processing steps. For example, even the best microscopes require computational alignment of tiles to form a mosaic, since mechanical tolerances and imperfect calibration introduce uncertainty into recorded tile positions. To enable assembly of a mosaic, tiles are slightly overlapped during acquisition so that each pair of adjacent tiles contains some identical cells. Image features in these cells are then used as reference points for “stitching” adjacent tiles into a seamless mosaic. In cyclic imaging (Gerdes et al., 2013; Lin et al., 2018), all tiles from the second and subsequent cycles must also be aligned to the mosaic through “registration” of image features across corresponding tiles. DNA-stained nuclei serve as an excellent image feature for alignment since they stain well with a variety of fluorescent dyes, are present at suitable density in most tissue types, and have sharp edges with high contrast. Multiple tools exist for registering image stacks and stitching image tiles (Chalfoun et al., 2017; Holtkamp and Goshtasby, 2009; Hörl et al., 2019) and some are available in common image analysis software such as ImageJ (Schneider et al., 2012). However, we have found that open source tools currently available for stitching and registering whole-slide images are unsatisfactory when applied to high-plex cyclic images with respect to speed, reliability and accuracy. Some commercial instruments have also integrated stitching routines, but we have found that these methods are only sufficient for visual review and are generally not accurate enough for quantitative single-cell analysis. Existing tools also struggle with very large images and generally require substantial format conversion and file renaming, a non-trivial task when confronted with 100 GB of data contained in 10^4^ megapixel-scale image tiles (a large 10-cycle whole-slide image).

In this paper we report the development of a new open source Python package, ASHLAR (Alignment by Simultaneous Harmonization of Layer/Adjacency Registration), for coordinated stitching and registration of multiplexed, multi-tile images. The package offers both a command line file-oriented interface and a documented API for incorporation into other tools. ASHLAR can directly process any image format supported by the widely used Open Microscopy Environment (OME) BioFormats library (Li et al., 2016) and it outputs standard OME-TIFF files. We describe ASHLAR’s design and implementation and compare its performance to existing tools using high-plex CyCIF images. ASHLAR is available as a Docker or Singularity container and has been incorporated into MCMICRO (Schapiro et al., 2022a), the Nextflow-based image processing pipeline developed by HTAN; as part of MCMICRO, ASHLAR has been tested with several hundred CyCIF, CODEX and mxIF images acquired from 12 types of mouse and human tissues at seven different institutions on five different microscopes and slide scanner platforms (**Supplementary Table 1**). ASHLAR is therefore a robust and practical tool for use with diverse spatial profiling methods.

## METHODS

### Overview of assembly process for cyclic multi-tile fluorescence images

ASHLAR operates in three broad phases to convert a multi-cycle multi-tile (Level 1) dataset into a cohesive (Level 2) mosaic image (Schapiro et al., 2022b) (**Figure 1**): (i) tiles within the first imaging cycle are stitched; (ii) tiles from the second and subsequent cycles are registered to corresponding tiles from the first cycle; and (iii) all tiles from all cycles are merged into a mosaic image. The output of stitching and registration is a list of new, corrected positions for all tiles in each cycle. Only in the final mosaic phase is the actual full-size many-channel mosaic image created. These mosaic images can be very large and contain information spanning length scales from less than 1 µm (subcellular structures) to cm in dimension (gross tissue morphology). In many cases, the boundary of a tissue specimen is irregular, and a significant fraction of the tiles in a rectangular data collection grid contain few if any cells, posing a challenge for stitching as well as an opportunity to reduce data collection demands by creating irregular-shaped tile sets that closely follow the outlines of the tissue.

**Figure 1:**
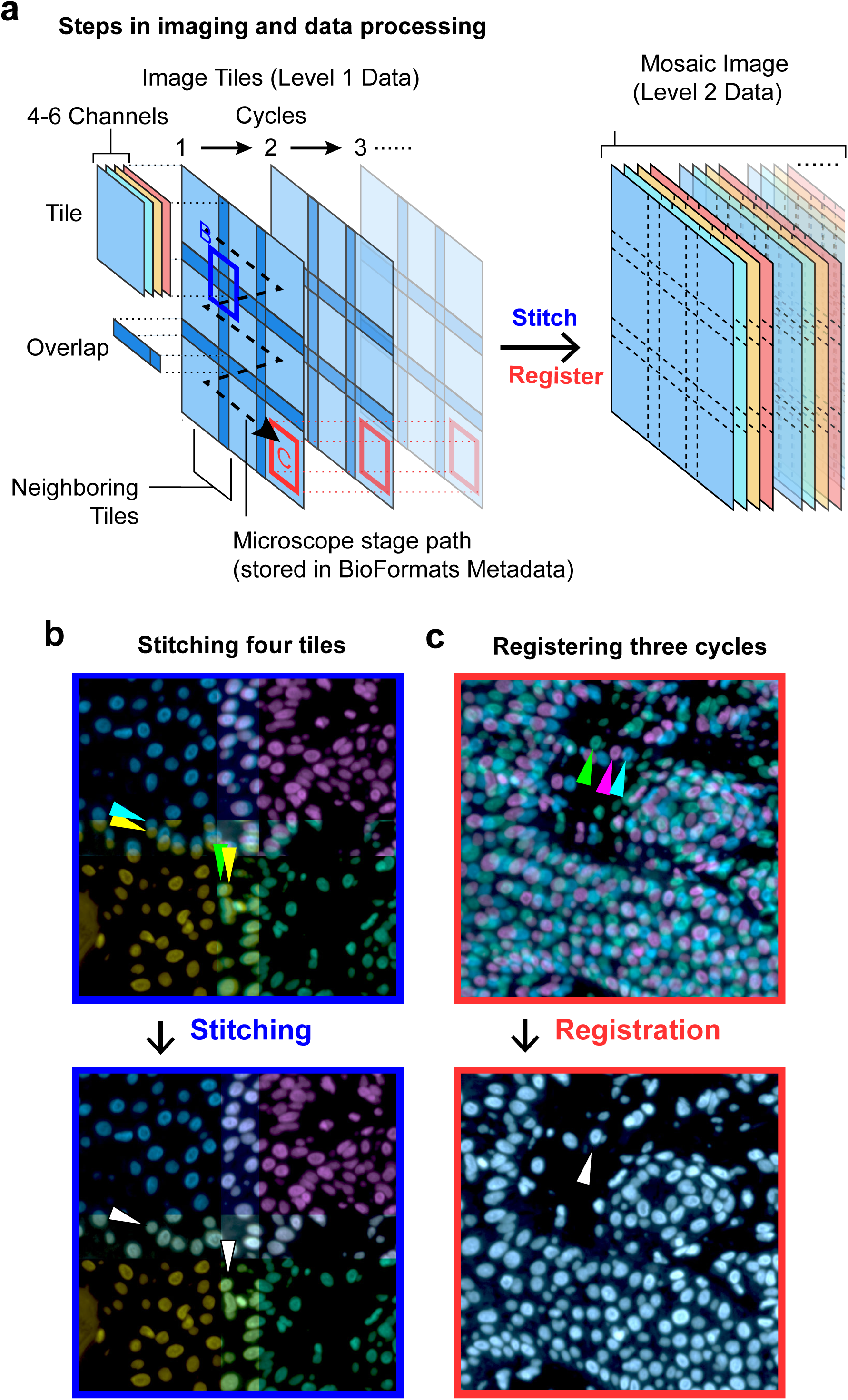
Schematic of cyclic whole-slide data acquisition, stitching, and registration. **(a)** One cycle of whole-slide imaging (scanning) is achieved by moving the microscope stage along a controlled path and acquiring multichannel image tiles that overlap. Further cycles repeat the process after the specimens are re-labeled with new antibodies or other detection reagents. Note that the left-hand portion of this panel depicts just a single reference channel (blue) across three cycles for clarity – actual data contains multiple channels and an arbitrary number of cycles. To integrate information across a wide spatial context at high resolution, it is necessary to stitch neighboring image tiles within one cycle and also register tiles across different cycles. **(b)** The corners of four neighboring tiles (Hoechst 33342-stained channel, pseudocolored by tile) from one cycle are positioned using the recorded microscope stage positions (upper panel) and the corrected stitched positions (lower panel). Arrows indicate two individual cells in the tile overlap regions before and after stitching. **(c)** The centers of three Hoechst-channel image tiles (pseudocolored by cycle) from different cycles are positioned using recorded stage positions (upper panel) and post-ASHLAR registered positions (lower panel). Arrows indicate one cell before and after registration.

To generate an initial estimate of tile positions, ASHLAR uses data from image tiles (grids of pixels), recorded stage positions, and physical pixel dimensions. ASHLAR leverages the Open Microscopy Environment Bio-Formats library (Li et al., 2016) to extract the necessary image data and metadata (stage position and pixel size) directly from native image files produced by the great majority of commercial microscopes, obviating the need for image format conversion and manual metadata extraction. For microscopes that do not support BioFormats, ASHLAR accepts a set of TIFF files using a configurable naming convention along with explicit specification of tile overlap and acquisition order. Subsequent stitching and registration involve aligning one image to another. In stitching, the small overlapping strips of adjacent tiles are aligned (**Figure 1b**), and in registration, full tiles that cover the same region of the sample but acquired in different cycles are aligned (**Figure 1c**). We performed stitching and registration only on the reference image channel (typically Hoechst 33342-stained nuclei) and applied the resulting positional corrections to all other channels recorded within that cycle. This is sufficient because the chromatic aberration exhibited by research-grade wide-field microscopes is not a major contributor to image inaccuracy at resolutions typically used for tissue imaging (10X to 40X magnification, 0.3 to 0.95 NA air objectives).

### Image alignment with sub-pixel precision phase correlation

ASHLAR uses the phase correlation algorithm (Kuglin and Hines, 1975) for image alignment during both stitching and registration phases. Phase correlation is a fast, parameter-free method that computes the image alignment with maximum cross-correlation, but it is only suitable for aligning images that are translated relative to each other in X and Y; it cannot directly align images that differ by rotation, scaling, skew, or non-affine transformations. This trade-off is acceptable for accurate stitching of multi-tile images on the five microscopes we have tested (see **Supplementary Table 2**) since the rigidity of the sample and the construction of modern stages ensures that almost all of the discrepancy between recorded stage positions and true positions can be modeled by translation alone. Stage position errors encountered between multiple imaging cycles are also purely translational, as long as the slide is always placed at exactly the same angle on the stage; this can routinely be achieved with kinematic mounts that positively register the slide in a consistent position (this is a standard feature of contemporary slide-scanning microscopes).

ASHLAR uses an enhanced method of phase correlation that improves the precision of tile alignment. The accuracy of classical phase correlation is limited to whole pixels. At pixel sizes around 1 µm or larger, this represents a substantial error relative to the size of a single cell. We overcame this limitation by using an improved phase correlation algorithm (Guizar-Sicairos et al., 2008) that offers arbitrary sub-pixel precision with minimal extra computation. An alignment precision of 0.1 pixels produced a discernible improvement in final mosaic quality over whole-pixel alignment, with diminishing returns beyond that. ASHLAR also enhances phase correlation by pre-filtering input images with the discrete Laplacian operator (or Laplacian of Gaussian operator – LoG – for noisy images) to eliminate auto-correlation. It has been understood for at least a century that computing cross-correlation can yield spurious results with signals that exhibit auto-correlation, but this fact is often overlooked in practice (Dean and Dunsmuir, 2016; Yule, 1926) – we are aware of only one open-source image stitching tool, ITKMontage (Zukić et al., 2021), that performs decorrelation. Our work with ASHLAR shows that decorrelation substantially improves confidence in image tile alignments.

### Image Stitching

The stitching procedure begins with the creation of a node-edge adjacency graph in which nodes represent tiles (**Figure 2** – step A1). Edges are added to the graph to connect overlapping tile pairs, which are initially identified by consulting recorded stage positions and other metadata. By reading recorded stage positions directly from BioFormats metadata, it is straightforward to support samples imaged with non-rectangular grids and irregular layouts. **Figure 3a** shows the overlap in adjacent tiles associated with one edge in the adjacency graph and reveals that one image is slightly translated relative to the next – this translation represents the stage positioning error we are trying to correct. When the overlap region contains many cell nuclei or other alignment features, phase correlation can accurately and confidently compute the correct translation between the images. Phase correlation will always return *some* value for translation of any two tiles, even when the overlap region is uninformative and contains only incidental signal or background noise in the registration channel; in these cases we rely on the recorded stage positions. Uninformative overlaps in tissue sections are most commonly encountered when nuclei are scant, such as in fat, connective tissue, or regions of necrosis, and in areas in which no tissue is present, such as along the edge of a specimen, between separate pieces of tissue, or between the circular cores that make up a tissue microarray (a regular grid of 50-200 0.3 to 1.2 mm pieces of tissue arrayed on a single microscope slide).

**Figure 2:**
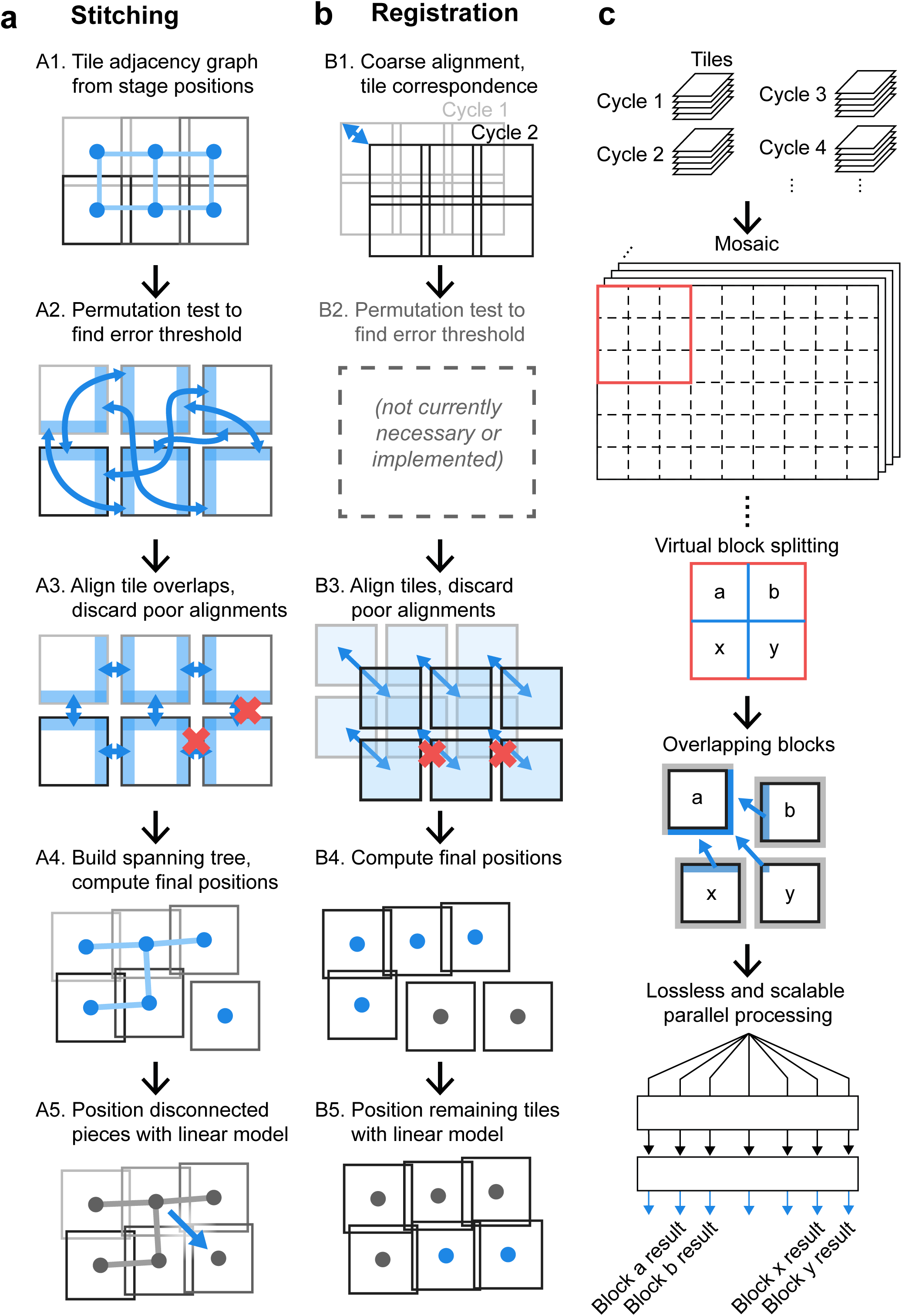
ASHLAR phases for aligning whole-slide scans. **(a)** Steps for stitching tiles within one cycle. **(b)** Steps for registering tiles across cycles. **(c)** Seamless mosaic generation enables whole-slide visualization and flexible re-tiling for downstream parallel processing. Blue-colored graphic components in each step depict the key elements or processes of that step. See text for details.

**Figure 3:**
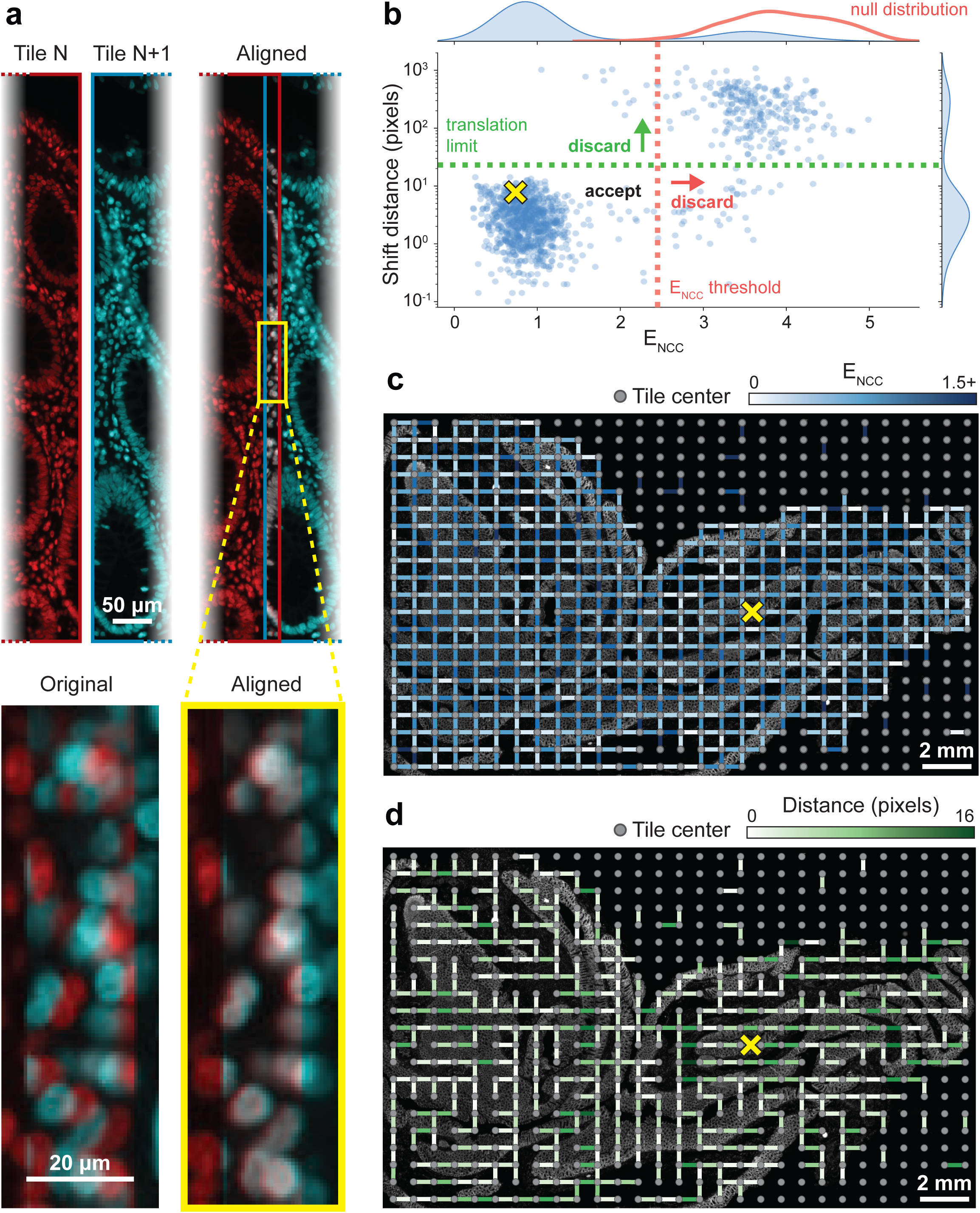
Visualizing stitching steps using a whole-slide scan from a colon specimen. All images and data in this figure derive from analysis of a large multi-tile image of human colon (see text for details). **(a)** Alignment of one pair of neighboring tiles from an image of human colon. Images of Hoechst 33342-stained nuclei in left and right tiles are pseudocolored in red and cyan respectively. The red and cyan images are overlaid before and after stitching to demonstrate the effect at the single-cell level. For context, in the remaining panels the location of this tile pair is denoted with a yellow X. **(b)** Alignment shift distance vs. E_NCC_ for all neighboring pairs, with E_NCC_ threshold and user-provided translation limit indicated. The null distribution generated by the permutation test (red) is overlaid on the E_NCC_ marginal distribution. Note that while the E_NCC_ threshold is computed as the 99^th^ percentile of the null distribution NCC values, it appears at the *left* end of the null distribution in this figure due to transformation of NCC to E_NCC_ by taking the negative logarithm (see text). **(c)** Adjacency graph with edges colored by E_NCC_ overlaid on the Hoechst image. Edges corresponding to discarded alignments (E_NCC_ or shift distance above the thresholds) are hidden. Hidden edges correlate with regions containing scant or no tissue. **(d)** Minimum spanning tree with edges colored by alignment shift distance.

We use normalized cross correlation (NCC) to score how well the translation returned by phase correlation aligns images, but the threshold dividing an effective alignment from a spurious one varies by dataset. We estimate this threshold by the 99^th^ percentile of NCC values computed from a permutation test that considers 1,000 randomly selected pairs of non-adjacent tiles (**Figure 2a** – step A2); this corresponds to the unadjusted one-sided empirical p-value threshold of 0.01. For each tile pair represented by an edge in the adjacency graph, we crop the images to their mutually overlapping region based on recorded stage positions and align them using phase correlation as described above. This yields a corrected X,Y shift, and NCC value (**Figure 2** – Step A3). For all downstream steps, we use the negative logarithm of the NCC values, hereafter referred to as E_NCC_ (NCC error), which provides a more intuitive “lower is better” error metric and empirically appears to have a more normal distribution. We use this E_NCC_ threshold and a user-provided translation limit parameter to determine whether to discard low-quality tile pair alignments (**Figure 3b**). The value of the translation limit is not particularly critical, as physical translation distances for spurious alignments tend to be fairly extreme (note that the Y-axis in Figure 3b is on a log scale). When a low-quality alignment is discarded, we delete the corresponding edge from the adjacency graph (**Figure 2** - step A3, **Figure 3c**). At this point, there are almost always more remaining pairwise alignments than tiles, leading to an overconstrained system. We solve this by constructing a minimum spanning tree with the E_NCC_ values as the edge weights and retaining only the alignments corresponding to edges in this tree (**Figure 2** **-** step A4). This allows us to discard extraneous alignments so the position of every tile is unambiguous. Since the edge deletion process in step A3 could split the graph into multiple disconnected pieces, we perform the spanning tree procedure independently for each piece.

With this spanning tree, it is straightforward to obtain final corrected positions by walking along the edges from tile to tile (starting at the root) and adding up each pairwise alignment along the way (**Figure 3d**). Even though individual pairwise tile alignments correct primarily for local uncorrelated stage position error, taken collectively they also characterize systematic errors such as miscalibrated pixel size or Z-axis camera rotation. To quantify these types of errors, we perform multiple linear regression of the corrected tile positions after stitching (dependent variable) against the original tile positions recorded by the microscope (independent variable) to generate a single affine transformation. This affine transformation is then used to adjust the relative positions of tiles with adjacency graphs that were split into multiple pieces (from step A3) to counteract systematic stage position error and improve accuracy (**Figure 2** **-** step A5). At the end of steps A1 to A5 (**Figure 2a**), optimized global positions have been determined for all tiles in the first cycle and, the stitching is complete.

### Image Registration

The procedure for registering subsequent cycles against the first cycle uses many of the same tools as stitching, although the goal is aligning whole tiles across data acquisition cycles rather than aligning adjacent tiles edge-to-edge within a single cycle (**Figure 2b**). First we establish a correspondence between each tile in the target cycle (the one to be registered) and the nearest tile in the first cycle by comparing recorded stage positions (**Figure 2b** - Step B1). Identifying these tile correspondences is trivial when recorded stage positions are consistent from run to run, and the geometry of the image acquisition grids is identical. However, a significant shift in stage positions can occur between cycles with microscopes that lack a physical “homing” procedure to zero stage position encoders at start-up. Shifts also arise where the planned tile grid is significantly displaced or rearranged between runs. To account for this shift when comparing tile positions, we down-sample the data by a factor of 20 and assemble low-resolution “thumbnail” mosaic images for each cycle using the recorded stage positions. We then align the thumbnails using phase correlation with sub-pixel precision to obtain a coarse alignment between the first and target cycles. Working with low-resolution images in this step saves compute time and memory while providing sufficient precision to accurately recover inter-cycle tile correspondences. Next, each corresponding tile pair is cropped to mutually overlapping regions and aligned using phase correlation (step B3). The resulting alignments are then filtered using the user-specified translation limit. Note that we do not currently use a permutation test and E_NCC_ threshold in the registration phase (step B2) as the translation limit alone has been sufficient for all images processed to date. For alignments that pass the filter, the target-cycle tiles are positioned by adding the alignment translations to the corrected positions of the corresponding tiles from the first cycle (step B4). The remaining tiles (generally those with sparse or no tissue, or where the tissue was damaged significantly during inter-cycle sample handling) are positioned using the affine transformation computed in the stitching phase, assuming that the same microscope and calibration conditions were used for both cycles (step B5). The registration steps described above are then applied to all other cycles, establishing corrected global positions for all remaining tiles. Importantly, our method registers each cycle of tiles against the first cycle (rather than consecutively against each preceding cycle) because each successive alignment step incurs additional error.

### Mosaic image generation

The result of the stitching and registration phases is a corrected global position for every image tile (**Figure 4**). To generate the final output image mosaic, we create an empty image large enough to encompass all corrected tile positions and copy each tile into it at the appropriate coordinates. Since each pairwise image registration is computed to a precision of 0.1 pixels as described above, the sum of these shifts for a given tile generally yields non-integer values for the final coordinates. ASHLAR defaults to applying sub-pixel translations on the tile images to account for this, but some users may prefer to round the final positions to the nearest pixel instead. Where neighboring image tiles overlap in the mosaic, they are combined with linear blending or one of several other user-selectable blending functions. The final many-channel image is then written out as a standard OME-TIFF file containing a multi-resolution image pyramid to support efficient visualization.

**Figure 4:**
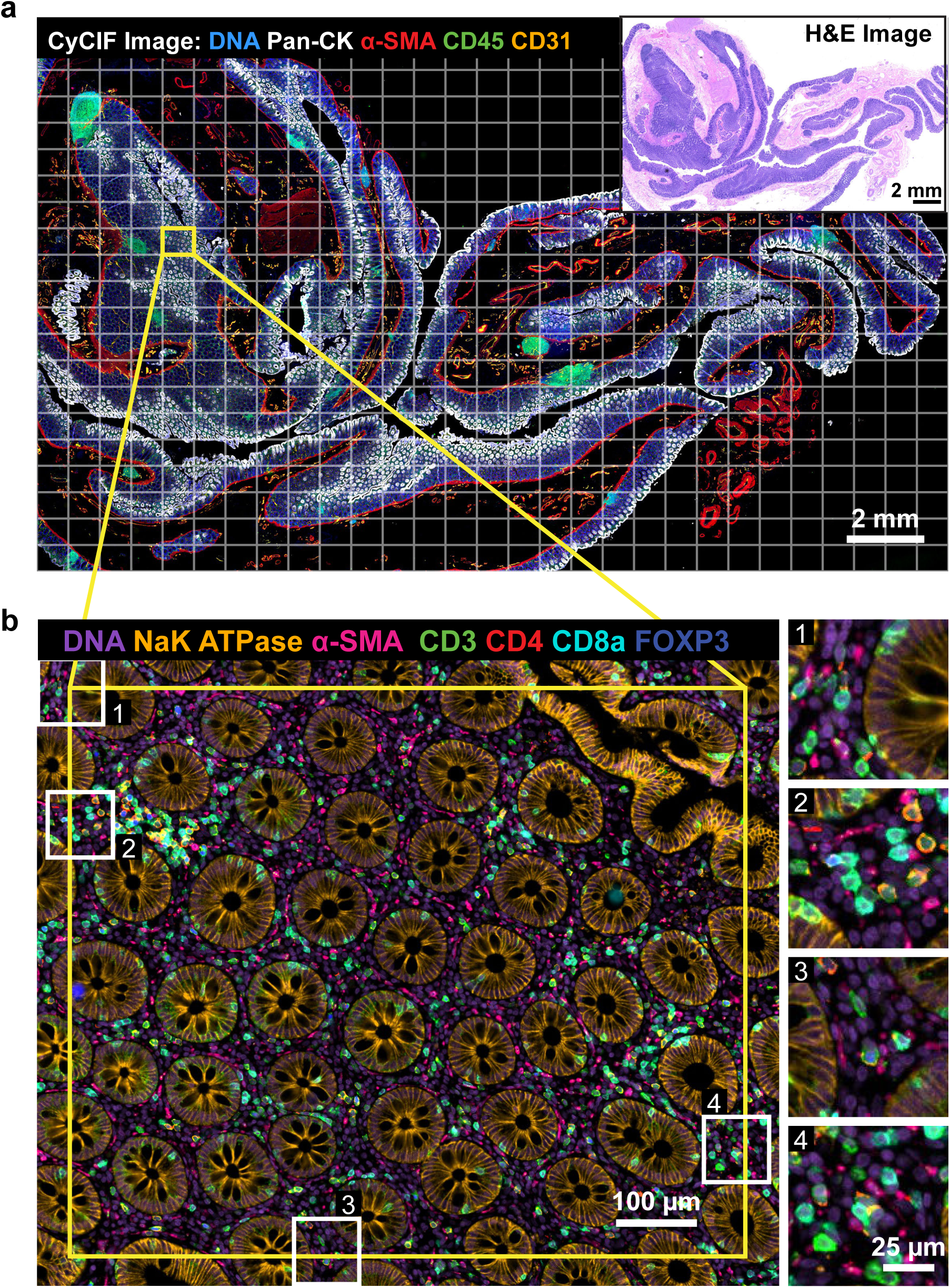
ASHLAR mosaic results. All images and data in this figure derive from analysis of the multi-tile image of human colon shown in Figure 3. **(a)** Pseudocolor image showing 5 channels from a 28-plex (9-cycle) t-CyCIF image of a normal human colon section acquired using the antibodies described in **Supplementary Table 3**. Tiles, denoted by the white grid, overlapped by ∼31 pixels (20 µm) **Inset**: Hematoxylin and eosin (H&E) staining of an adjacent section of the same specimen. **(b)** Higher magnification view of the area surrounding a single tile showing 7 channels from 4 different cycles to highlight stitching and registration accuracy. Insets 1-4 depict regions of the tile overlap areas at full resolution (note that the antibodies shown in panels **a** and **b** differ to make structures relevant to different spatial scales more apparent).

### Implementation

ASHLAR is implemented in Python 3 and utilizes many numerical and image processing routines from the numpy, scipy, scikit-image, scikit-learn and networkx packages. The pyjnius Python-to-Java connector provides access to the Java BioFormats library for reading microscopy image files. The user interface is mainly via command-line script but the underlying Python modules may also be used directly.

## RESULTS

### Evaluation of stitching accuracy

We identified MIST (Chalfoun et al., 2017) as the current state-of-the-art public-domain tool for stitching large, tiled microscopy images. We used the evaluation framework described by Chalfoun et al. to compare the accuracy of stitching by ASHLAR and MIST using that manuscript’s most challenging dataset: a plate of widely-spaced GFP-labeled stem cell colonies that were grown for 2 days and imaged with 10% tile overlap. The Chalfoun et al. dataset contains two image sets acquired via separate mechanisms: i) images acquired with “traditional” overlapping tiles and ii) ground-truth images – with each colony centered and wholly contained in a single image field – acquired with a closed-loop microscope stage control algorithm. The Chalfoun et al. evaluation framework assesses the accuracy of a stitching algorithm by applying the algorithm to the overlapping tile set, segmenting the stitched output mosaic into colonies, and finally comparing each colony’s area and position to the ground truth data. Four metrics are reported: *False negative count* (FN), *false positive count* (FP), per-colony *size error* (S_err_), and per-colony position *distance error* (D_err_) (see Chalfoun et al. for full details). MIST and ASHLAR yielded the same false negative and false positive counts (FN = 47, FP = 4), the size error distributions were nearly the same and had indistinguishable medians (median S_err_ = 0.0474%), and the distance error distributions were also similar with a difference between medians that was not statistically significant (MIST median D_err_ = 10.8 pixels, ASHLAR median D_err_ = 11.5 pixels, Mood’s median test p-value = 0.32) (**Figure 5a**). We conclude that MIST and ASHLAR have similar stitching accuracy when applied to a previously described test set involving cells grown *in vitro*.

**Figure 5:**
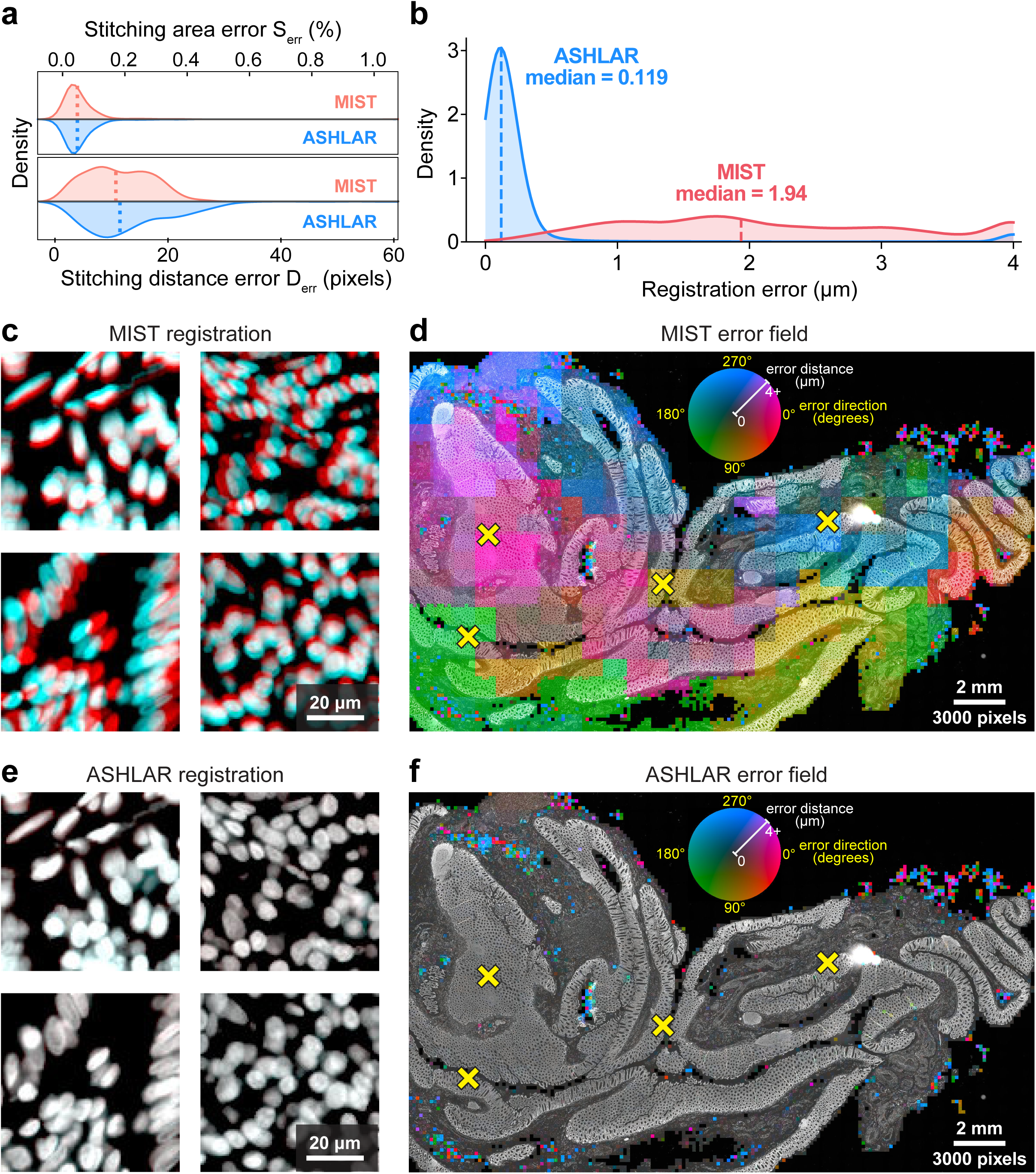
Performance comparison of ASHLAR and MIST software. **(a)** Stitching error metric distributions (kernel density estimate) for MIST and ASHLAR computed according to the stitching evaluation framework of Chalfoun et al. (Chalfoun et al., 2017). Dotted lines indicate median values; neither difference in medians was statistically significant. **(b-f)** Images and data derive from analysis of the multi-tile image of human colon shown in Figures 3-4. **(b)** Local registration error distance distributions for MIST and ASHLAR mosaic images of two t-CyCIF cycles of a human colon section. Distances at the upper end in this plot as well as in panels d and f were clipped to the 90^th^ percentile of the MIST error values (∼4 µm) to highlight the relevant data. **(c)** Full-resolution view of four regions from the MIST mosaics demonstrating local registration error in different directions. The Hoechst images of nuclei from cycles 1 and 2 are pseudocolored red and cyan, respectively, to visualize the effect of registration error at the single-cell level. The MIST median error of ∼2 µm is around one quarter of the diameter of the average cell nucleus, a shift that is clearly visible at full resolution. **(d)** Heatmap of MIST local registration error direction (hue) and magnitude (intensity) at 200-pixel resolution overlaid on the Hoechst image (brighter colors indicate larger errors). Characteristic tile-sized scale of heatmap features suggests inconsistent stitching. Yellow X marks indicate locations highlighted in panel c. **(e)** The same regions as in panel c, but taken from ASHLAR mosaics. An identical pseudocoloring scheme is used; the red and cyan images, now more accurately registered, combine to appear nearly white. The ASHLAR median registration error of ∼0.1 µm is approximately 1% of the diameter of a nucleus. **(f)** Heatmap of Ashlar local registration error using the same intensity and hue scale as in panel d showing overall lower error and no apparent tile-scale features. Remaining small-scale errors represent damaged tissue that could not be registered.

### Benchmark dataset for evaluating registration accuracy

As a first test of ASHLAR and MIST on high-plex, whole-slide tissue data, we acquired a ∼24 mm x 14 mm x 5 µm thick section of a human normal colon sample from the Cooperative Human Tumor Network (https://www.chtn.org/). This sample was subjected to 9-cycle t-CyCIF (Lin et al., 2018) to generate a subcellular-resolution 28-plex image. Cell nuclei were stained with Hoechst 33342 in every cycle, providing reference features for image alignment. Imaging was performed on a RareCyte CyteFinder II HT Instrument with a 20X 0.75 numerical aperture (NA) objective and four excitation and emission filter pairs having peak and full-width at half-maximum bandpass wavelengths (in nm) of: 395/25-438/26, 485/25-522/20, 555/20-590/20, and 651/10-692/40, respectively. 2 x 2 pixel binning was used during acquisition yielding a 4-channel tile with dimensions of 1280 x 1080 pixels and a pixel size of 0.65 µm per pixel. To image the entire specimen, a grid of 609 (29 x 21) tiles was used and each tile overlapped by ∼31 pixels (20 µm or 2-3%) in both directions. Each cycle yielded one OME BioFormats-compatible RCPNL file containing 16-bit imaging data from all 609 tiles, approximately 7 GB in size. The entire 9-cycle dataset comprises 5,481 image tiles and is 61 GB in size. The experimental protocol is documented on protocols.io (https://dx.doi.org/10.17504/protocols.io.bjiukkew) and antibodies used are listed in **Supplementary Table 3**. The primary unstitched image data are freely available for download from Synapse at https://dx.doi.org/10.7303/syn25826362. We ran both MIST and ASHLAR on this colon dataset, yielding mosaic images approximately 36,000 x 22,000 pixels in dimension. **Figure 4a** shows the resulting image mosaic following stitching and registration with ASHLAR; the quality of the alignment is highlighted at four regions of tile-tile overlap in **Figure 4b**. We then used the Hoechst reference channel mosaic images from the first two cycles to evaluate whether the stitched and registered output from MIST and ASHLAR was aligned accurately enough for single-cell-level quantitative analysis.

### Image registration of independently stitched mosaics

Whereas ASHLAR performs stitching and registration in a coordinated process, it was necessary to globally register the MIST output mosaics before evaluating local registration accuracy. We first downsampled the MIST mosaic images by 10X to obtain a manageable image size and then aligned them with subpixel-precision phase correlation (phase correlation on the full-size images required a computer with more RAM than was readily available to us). We performed the same process on the ASHLAR mosaics to verify that their global registration was already optimal.

### Optical flow computation and evaluation of registration accuracy and robustness

We used dense optical flow fields to quantify and visualize local registration errors in the Hoechst reference channel. Because we could not find any general-purpose dense optical flow implementations capable of processing gigapixel-scale images on a reasonable workstation computer, we implemented our own approximate method suitable for small-magnitude flow fields using a block-based approach which is memory-efficient and highly parallelizable. The two images to be compared are broken down into non-overlapping blocks of 200 x 200 pixels and the relative shift between each corresponding pair of blocks is computed using phase correlation. Any minor rotation, scaling, or shear between the full input images is then accounted for through a compensating affine transformation computed via multiple linear regression on the full set of per-block shift vectors. This phase correlation and transformation procedure yields a 200x-downsampled “block-dense” flow vector field that characterizes the local registration error. It is important to note that there is no separate ground truth data in this method – it only measures the consistency of a stitching/registration algorithm against itself. Having previously established that Ashlar and MIST have equivalent stitching accuracy, we felt this approach was reasonable.

We defined the local registration error as the magnitude of the flow vector field at each point. The median error was 1.94 µm for the MIST mosaics (∼3 pixels) and 0.119 µm (∼0.2 pixels) for the corresponding ASHLAR mosaics (**Figure 5b**). At a magnification sufficient to see individual cells, the error generated by MIST was readily apparent (**Figure 5c**). Visualizing the full vector field on top of the reference channel images (**Figure 5d**) showed that the field direction was often consistent across large regions but could change dramatically at tile boundaries. This most likely arises because small local stitching differences propagate across the mosaic in a manner that is uncorrelated between cycles, leading to inter-cycle shifts that cannot be corrected by any rigid adjustment of the entire mosaic. It is important to note that this is not a weakness of MIST *per se,* but rather a consequence of applying a tool designed for stitching alone to the combined process of stitching and registration, a use case for which MIST was not designed. With the ASHLAR-generated mosaic, vector field visualizations confirmed a much lower level of registration error (**Figure 5e, f**). Close inspection of the few regions with high error showed that they corresponded to parts of the tissue in which cells had become physically distorted or detached from the slide as a consequence of the cyclic staining procedure. Thus, remaining errors are not a result of errors in registration and stitching but rather extrinsic processes that must be identified and accounted for by downstream error-checking procedures.

To demonstrate ASHLAR’s robustness and versatility with different image types, we compared its registration accuracy against MIST on four further datasets encompassing two cyclic imaging techniques, three vendors’ microscopes, and two new tissue types plus a tissue microarray (TMA) that itself spans a multitude of tissue types and disease states. We also evaluated one additional open-source stitching algorithm, BigStitcher (Hörl et al., 2019), on some of these datasets. The datasets are described in **Supplementary Table 4** and the evaluation results shown in **Supplementary Figures S1-S5**. Overall, ASHLAR compared favorably to MIST and BigStitcher on all datasets with respect to registration of cyclic datasets.

### Evaluation of commercial stitching algorithms

Slide scanning microscopes include stitching algorithms as part of their data acquisition and analysis software. These algorithms use proprietary file types and in most cases they erase the original image tiles or strips after generating the final stitched output image. Thus, detailed evaluation of their performance is not straightforward, but it is possible to evaluate their stitching consistency by using our optical flow method to examine two re-scans of the same specimen. We performed such an analysis using a human colorectal adenocarcinoma specimen retrieved from the archives of the Department of Pathology at Brigham and Women’s Hospital with Institutional Review Board (IRB) approval as part of a discarded/excess tissue protocol. The specimen was stained with H&E and scanned in brightfield mode at the Brigham and Women’s Hospital Pathology Core Facility using three different slide scanning microscopes: a Leica Aperio GT450, Leica Aperio VERSA, and Hamamatsu NanoZoomer 2.0-HT. We also imaged the adenocarcinoma specimen in brightfield mode using a GE INCell 6000 microscope to produce tiles suitable for processing with ASHLAR. The specimen was imaged twice on each instrument to emulate a cyclic imaging workflow. Pre-stitched mosaic image pairs generated by the three scanners and the ASHLAR-stitched mosaic pair assembled from the INCell 6000 tiles were subjected to the registration accuracy evaluation described above. The results, shown in **Figure 6**, demonstrate that all of the tested systems exhibit worse stitching consistency than ASHLAR. Inspection of the underlying images reveals obvious stitching errors that would confound single-cell-level analysis. We also observed that the error field images also contained “signatures” of each instrument’s internal design, such as line sensor vs. area sensor and sensor size and orientation. Thus, commercial algorithms included with existing slide scanners do not appear to fully correct for intrinsic limitations of the instrumentation. Finally, we evaluated the registration accuracy of the stitching feature in Zeiss’s Zen software which was recently used to generate a publicly available 50-plex rat brain dataset (Maric et al., 2021) based on cyclic fluorescence imaging on a Zeiss Axio Imager Z2 microscope. We compared the DNA channel images from two imaging cycles from this dataset with our evaluation framework and found conspicuous errors here as well (**Supplementary Figure S6)**. Thus, while commercial algorithms may stitch well enough for visual review and gross structural analysis, they have weaknesses that are very likely to impact single-cell quantification – especially with cyclic imaging methods.

**Figure 6:**
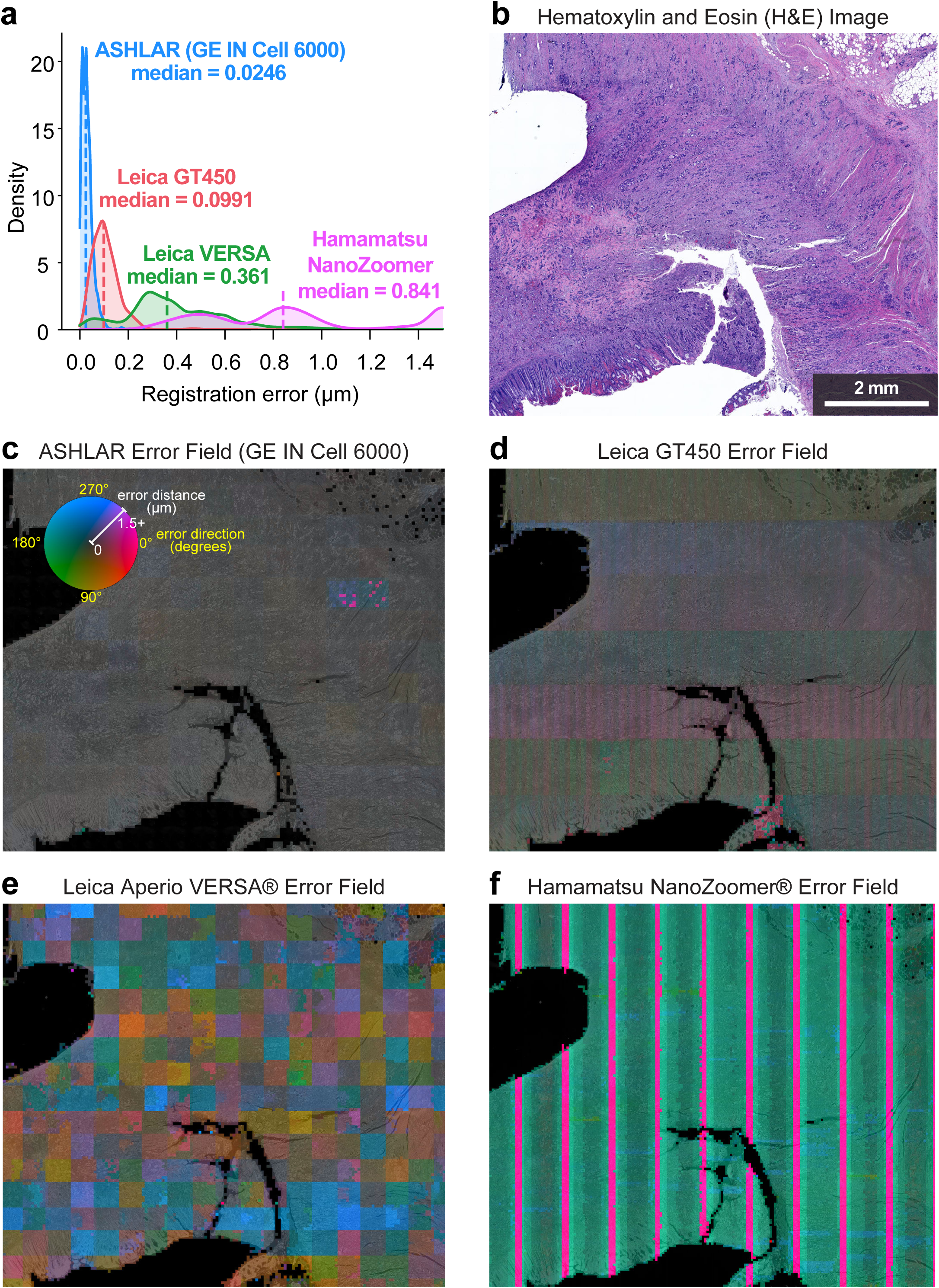
Comparison of registration accuracy between ASHLAR and software included with various commercial slide scanners. All images and data in this figure derive from a single section of a human colorectal adenocarcinoma biopsy (see text for details). **(a)** Local registration error distance distributions for technical replicate slide scans on three dedicated slide scanning microscopes as well as an ASHLAR mosaic from a research-grade microscope. All scans used the same H&E-stained section of a human colon adenocarcinoma biopsy to allow direct comparison of results. Distances in this panel as well as panels c-f were clipped to 1.5 µm at the upper end to highlight the relevant data. **(b)** H&E staining of the sample used for this analysis. **(c)** Heatmap of ASHLAR local registration error direction (hue) and magnitude (intensity) at 200-pixel resolution overlaid on the blue channel of the brightfield image, inverted (bright becomes dark and vice versa). **(d-f)** Heatmap of the three slide scanners’ local registration error, as in panel c. Engineering details of the different instruments are readily apparent in the error field patterns.

### Runtime and memory usage

The runtime and RAM usage for the stitching and registration phases of ASHLAR are each comparable to that of MIST stitching running on a single CPU core. On the first two cycles of the human colon dataset, ASHLAR required 306 seconds (149 s to stitch cycle 1 and 157 s to register cycle 2) and 2.1 GB RAM. MIST-FFTW required 228 seconds (114 s per cycle) and 2.5 GB RAM to compute corrected tile positions. If we include the 30 seconds per cycle required to convert the datasets from the microscope vendor’s native file format into MIST’s required single-TIFF format (ASHLAR requires no such conversion step) the total for MIST is 288 seconds. MIST does execute more quickly when allowed to use multiple CPU cores or a GPU; we have not enabled parallel processing in ASHLAR but expect a similar increase in speed. ASHLAR’s runtime per imaging cycle varies linearly with the total number of pixels in all tiles of the reference channel (**Supplementary Figure S7)**. All measurements were taken on a 3.5 GHz Intel Xeon E5-1620 v3 CPU with 32 GB of RAM and a SK Hynix SH920 512 GB SSD running Ubuntu Linux 20.04. Software versions were as follows: ASHLAR 1.14.0, MIST 2.0.7, Python 3.8.10, OpenJDK 1.8.0.

## DISCUSSION

To date, ASHLAR has been used to stitch several hundred whole-slide images collected using diverse acquisition technologies and instruments (**Supplementary Table 1**). ASHLAR, therefore, provides a robust and efficient way to generate large, multi-channel, mosaic images of tissues and other biological specimens by assembling individual megapixel image tiles collected at multiple wavelengths over multiple imaging cycles. The key innovation for image quality is joint optimization of stitching and registration as opposed to stitching individual cycles independently and then attempting to register mosaic images against each other. Joint optimization becomes increasingly important as the size of the specimen increases. Coupling ASHLAR with tile-based image acquisition and cyclic data collection makes it possible to optimally balance the resolution, size, and plex of a tissue image for reliable analysis of spatial features on a wide variety of scales. Although many recent highly multiplexed studies have relied on small fields of view and TMAs, whole-slide imaging is emerging not only as a diagnostic necessity (Health, 2019) but also as a key requirement for basic research into the spatial organization of tissue and tumor microenvironments (Lin et al., 2021). ASHLAR is optimized for these data acquisition requirements and is more rapid and accurate than existing open-source methods we have tested as well as commercial software available with many slide scanners. ASHLAR reads and writes files in the OME-TIFF standard and can process images from almost all commercial microscopes using the OME Bio-Formats library (Li et al., 2016). This greatly streamlines the stitching and registration process since little manual intervention is required. Once an optimized whole-slide image mosaic has been generated, it is often convenient to visualize or analyze limited regions of the image. ASHLAR, therefore, supports re-tiling using adjustable block sizes and overlaps while retaining subpixel registration and without losing any information along the original tile seams (**Figure 2c**). This block-based processing is critical for downstream image processing such as single-cell quantification and pixel-level machine learning, as few methods can process gigapixel-scale images in a single pass.

ASHLAR was designed as a general-purpose algorithm compatible with a wide variety of microscopes and image acquisition technologies. To establish that ASHLAR meets these requirements, we incorporated it into MCMICRO (Schapiro et al., 2022a), a data processing pipeline leveraging either Docker or Singularity containers (Kurtzer et al., 2017; Merkel, 2014) and implemented it in the workflow systems Nextflow (Di Tommaso et al., 2017) and Galaxy (Afgan et al., 2018). MCMICRO makes it possible to process high-plex tissue images of raw data into a table of computed single-cell features; stitching and registration by ASHLAR is an essential early step in the MCMICRO pipeline. Via MCMICRO, ASHLAR has been made available to multiple research teams in the NCI HTAN consortium (Rozenblatt-Rosen et al., 2020) and, to date, has been used successfully in 17 published manuscripts and three posted pre-prints by investigators at seven different institutions. These papers encompass a total of ∼240 whole-slide images and 11 TMAs that have been successfully stitched using data obtained with three data acquisition methods (CyCIF, CODEX, and mIHC (Gerdes et al., 2013; Goltsev et al., 2018; Lin et al., 2018; Tsujikawa et al., 2017)) and on five different types of microscopes (**Supplementary Table 1**). This experience demonstrates that ASHLAR operates as designed with most image data, but some edge cases may require tuning the algorithm’s parameters. This is most commonly encountered when tile overlaps are too small or the tissue has suffered grievous damage during processing. The online documentation for ASHLAR (available at https://labsyspharm.github.io/ashlar/) discusses these and related issues in greater detail. ASHLAR is available under the permissive MIT open source software license, making it possible for commercial companies to modify and package it with their instruments.

ASHLAR is effective not only with conventional rectangular image acquisition grids, but also with images involving multiple non-overlapping regions of interest and tiles that do not form a rectangular grid. The ability to manage irregular and disconnected specimens has emerged as a key requirement in the broader application of tissue imaging. By instructing a microscope to avoid imaging empty space lying outside of the margins of the tissue, imaging time and file size can be reduced, often by a factor of two or more (a significant advantage as datasets approach terabyte scale). We have successfully used ASHLAR to assemble images of tissue microarrays (TMAs), in which 0.3 to 1 mm diameter “cores” from multiple tissue specimens are positioned in a regular array on a glass slide, making it possible to analyze over 100 specimens in parallel. This represents a potentially challenging stitching problem since much of the slide is devoid of sample, and individual cores are often divided among multiple fields. Core biopsies and fine needle aspirations are other samples in which collection of non-rectangular images is highly advantageous. Such biopsies are typically long and thin (0.3 x 10-20 mm) and rarely aligned along the axis of the slide, making it necessary to collect tiles on a diagonal. The ability of ASHLAR to reject spurious alignments using a permutation test followed by pruning of adjacency graphs make the algorithm robust to regions of the images such as these that contain little if any data in the registration channel.

Much of the recent discussion about multiplexed tissue imaging has focused on the importance of the number of channels (assay plex) (Baharlou et al., 2019), since more channels allow more proteins or genes to be assayed and yield more detailed molecular insights. However, two other considerations are at least as important: image resolution and field of view (speed also matters for high volume applications). In the case of fluorescent imaging of a single tile, resolution and field of view are functions of the optics, primarily the numerical aperture of the objective lens, the properties of the transfer optics and the number of camera pixels – which together specify pixel size (Ghiran, 2011). For large whole slide images assembled from many tiles, the accuracy of image stitching and registration also becomes critical. ASHLAR directly addresses this requirement. In most applications, it is also combined with other software to optimize the quality of image mosaics. Prior to stitching and registering tiles using ASHLAR we perform illumination correction using BaSiC (Peng et al., 2017), which exploits low-rank and sparse decomposition to correct for uneven shading and background variation in microscope images. This is essential because the illumination of each tile is typically brightest at the center of the field (along the optical axis) and dimmer at the edges.

### Limitations

We have found that the spanning tree approach used to combine individual pairwise tile alignments is broadly effective. However, one recurrent weakness observed with large specimens is that tiles at the margin of the tissue that are adjacent in physical space are often found to lie far apart in the adjacency graph. When corrected positions are determined, uncorrelated error accumulates as pairwise shifts are added up along the edges in the spanning tree. The resulting stitching error is rarely noticeable by eye in the resultant mosaic image, but we have identified this as an area for future improvement of the algorithm.

To achieve reasonable processing speed, ASHLAR makes some compromises with respect to the factors it accounts for during stitching and registration. For example, ASHLAR currently performs only rectilinear correction of tile position and assumes tiles have the same magnification. When images from different microscopes must be combined it is usually necessary to account for changes in camera angle due to rotation of individual cameras and their microscope stages relative to each other. Scaling is also frequently different across instruments, even when the same objective is used, due to differences in transfer optics and sensor configurations. We have never encountered the need to assemble an image from multiple microscopes (partly because many other batch effects are introduced) but corrections for image rotation and scaling can be performed through minor additions to the alignment procedure (Gonzalez, 2011); we will add these features to ASHLAR as the need arises, most likely arising from a requirement to combine multiple different imaging modalities.

## ACKNOWLEDGEMENTS

We thank Jerry Lin, Zoltan Maliga, Connor Jacobson, Artem Sokolov and other members of the Laboratory of Systems Pharmacology for their assistance. The normal colon specimen used for this work was acquired from the Cooperative Human Tumor Network (https://www.chtn.org/). We thank the members of the HTAN Consortium for their support; a full list can be found at humantumoratlas.org.

## Data availability

The human normal colon images used to evaluate registration accuracy are available in Synapse at https://dx.doi.org/10.7303/syn25826362 (this sample has been judged to be “not human subjects research”). This and all other data used is described in **Supplementary Table 4**.

