## Supplementary Table 1 for "Stitching and registering highly multiplexed whole slide images of tissues and tumors using ASHLAR"

### SUPPLEMENTARY TABLE 1. Published image data stitched and registered using ASHLAR.

| Authors | Year | DOI | Tissue type | Tissue or organ of origin | WSI | TMA |
|---|---|---|---|---|---|---|
| Schapiro et al | 2021 | doi.org/10.1038/s41592-021-01308-y | Normal tonsil, colorectal cancer | Tonsil (2), colon (3), TMA (2) | 5 | 2 |
| Rashid et al | 2021 | doi.org/10.1038/s41551-021-00789-8 | Lung carcinoma | Lung (1), | 1 | 0 |
| Duraiswamy et al | 2021 | doi.org/10.1016/j.ccell.2021.10.008 | Ovarian cancer | Ovarian (19) | 19 | 0 |
| Krueger et al | 2021 | doi.org/10.1109/TVCG.2021.3114786 | Colorectal cancer, lung adenocarcinoma | Colon (2), lung (1) | 3 | 0 |
| Gaglia et al | 2022 | doi.org/10.1101/2021.05.16.443704 | Breast carcinoma, ovarian carcinoma | Breast (4), ovarian (0), TMA (6) | 4 | 6 |
| Liu et al | 2021 | doi.org/10.1038/s41591-021-01331-8 | Melanoma | Skin (32) | 32 | 0 |
| Keenan et al | 2021 | dx.doi.org/10.1158%2F1078-0432.CCR-20-3089 | Breast carcinoma | Breast (12) | 12 | 0 |
| Hemming et al | 2021 | dx.doi.org/10.1158%2F1078-0432.CCR-20-3538 | Gastrointestinal stromal tumor | Gastrointestinal tract | 0 | 1 |
| Mehta et al | 2021 | doi.org/10.1038/s43018-020-00148-7 | Breast carcinoma | Breast (16) | 16 | 0 |
| Iorgulescu et al | 2021 | doi.org/10.1158/1078-0432.CCR-20-2291 | Glioblastoma | Brain | 2 | 0 |
| Ringel et al | 2021 | doi.org/10.1016/j.cell.2020.11.009 | Colorectal tumors | Colon (14) | 14 | 0 |
| Chandrashekar et al | 2020 | doi.org/10.1126/science.abc4776 | Lung | Lung (2) | 2 | 0 |
| Hemming et al | 2020 | doi.org/10.1200/po.19.00287 | Sarcoma | Liver (2), sacrum (1) | 3 | 0 |
| Färkkilä et al | 2020 | doi.org/10.1038/s41467-020-15315-8 | Ovarian cancer | Ovarian (19) | 19 | 0 |
| Gaglia et al | 2020 | doi.org/10.1038/s41556-019-0458-3 | Colorectal tumors | Colon (1) | 1 | 0 |
| Krueger et al | 2019 | doi.org/10.1109/TVCG.2019.2934547 | Melanoma, breast carcinoma, lung adenocarcinoma | Skin (1), breast (1), lung (1) | 3 | 0 |
| Du et al | 2019 | doi.org/10.1038/s41596-019-0206-y | Normal tonsil, lung carcinoma | Tonsil (1), lung (1), lymph node (1), brain (1) | 4 | 0 |
| Nirmal et al | 2022 | doi.org/10.1101/2021.05.23.445310 | Melanoma | Skin (22) | 22 | 0 |
| Lin et al | 2022 | doi.org/10.1101/2021.03.31.437984 | Colorectal cancer | Colon (75), TMA (2) | 75 | 2 |
| Kalocsay et al | 2021 | doi.org/10.1101/2020.10.14.339952 | Lung | Lung (1) | 1 | 0 |
| **Total** | | | **12 tumor/tissue types** | | **238** | **11** |
