## Supplementary Table 2 for "Stitching and registering highly multiplexed whole slide images of tissues and tumors using ASHLAR"

**SUPPLEMENTARY TABLE 2. Microscopes Tested with ASHLAR.**

| Instrument | Type | Objective | Field of View | Nominal Resolution[1] |
|---|---|---|---|---|
| RareCyte CyteFinder | Slide scanner | 10X/0.3NA | 1.66 x 1.40 mm | 1.06 µm |
| | | 20X/0.75NA | 0.83 x 0.7 mm | 0.42 µm |
| | | 40X/0.6NA | 0.42 x 0.35 mm | 0.53 µm |
| RareCyte Orion | Slide scanner | 20X/0.75NA | 0.66 x 0.66 mm | 0.42 µm |
| | | 40X/0.95NA | 0.33 x 0.33 mm | 0.33 µm |
| GE IN Cell Analyzer 6000 | Slide scanning mode | 10X/0.45NA | 1.3 x 1.3 mm | 0.70 µm |
| | | 20X/0.75NA | 0.66 x 0.66 mm | 0.42 µm |
| | | 40X/0.95NA | 0.33 x 0.33 mm | 0.33 µm |
| | | 60X/0.95NA | 0.22 x 0.22 mm | 0.33 µm |
| GE IN Cell Analyzer 6000 | Confocal mode | 60X/0.95NA | 0.22 x 0.22 mm | 0.21 µm |
| Zeiss Axio Scan.Z1 | Slide scanner | 10X/0.45NA | 1.3 x 1.3 mm | 0.70 µm |
| | | 20X/0.8NA | 0.66 x 0.66 mm | 0.40 µm |
| Zeiss Axio Observer.Z1 | Slide scanner | 20X/0.8NA | 0.83 x 0.66 mm | 0.40 µm |

[1]The nominal resolution (r) was determined using the formulae: $(r) = 0.61\lambda/NA$ for widefield or $(r) = 0.4\lambda/NA$ for confocal microscopy ($\lambda = 520$ nm). The actual resolution depends on optical properties, the thickness of the tissue section, and both the alignment and the quality of the optical components used.
