## Supplementary Table 3 for "Stitching and registering highly multiplexed whole slide images of tissues and tumors using ASHLAR"

**SUPPLEMENTARY TABLE 3. Antibodies used for CyCIF imaging of the human colon specimen.**

| Cycle Number | Channel Number | Target Name | RRID identifier | Fluorophore | Clone | Vendor | Catalog Number |
|---|---|---|---|---|---|---|---|
| 1 | 1 | DNA | AB_10626776 | Hoechst 33342 | - | Cell Signaling Technology | 4082S |
| 1 | 2 | Rabbit-IgG | AB_2534114 | Alexa Fluor 488 | - | Invitrogen | A-11070 |
| 1 | 3 | Rat-IgG | AB_2535855 | Alexa Fluor 555 | - | Invitrogen | A-21434 |
| 1 | 4 | Mouse-IgG | AB_2535806 | Alexa Fluor 647 | - | Invitrogen | A-21237 |
| 2 | 5 | DNA | AB_10626776 | Hoechst 33342 | - | Cell Signaling Technology | 4082S |
| 2 | 6 | Na/K ATPase | AB_2798866 | - | D4Y7E | Cell Signaling Technology | 23565 |
| 2 | 7 | CD3 | AB_2889189 | - | CD3-12 | Abcam | ab11089 |
| 2 | 8 | Cellular tumor antigen p53 | AB_2206626 | - | DO7 | Dako | M7001 |
| 3 | 9 | DNA | AB_10626776 | Hoechst 33342 | - | Cell Signaling Technology | 4082S |
| 3 | 10 | Antigen Ki67 | AB_2687824 | Alexa Fluor 488 | D3B5 | Cell Signaling Technology | 11882 |
| 3 | 11 | Pan-cytokeratin | AB_11217482 | eFluor 570 | AE1/AE3 | eBioscience/Thermo Fisher | 41-9003-80 |
| 3 | 12 | Aortic smooth muscle actin | AB_2574361 | eFluor 660 | 1A4 | eBioscience/Thermo Fisher | 50-9760-80 |
| 4 | 13 | DNA | AB_10626776 | Hoechst 33342 | - | Cell Signaling Technology | 4082S |
| 4 | 14 | CD8a | AB_2574412 | Alexa Fluor 488 | AMC908 | eBioscience/Thermo Fisher | 53-0008-80 |
| 4 | 15 | CD4 | AB_2573601 | eFluor 570 | N1UG0 | eBioscience/Thermo Fisher | 41-2444-80 |
| 4 | 16 | CD45 | AB_493034 | Alexa Fluor 647 | HI30 | BioLegend | 304020 |
| 5 | 17 | DNA | AB_10626776 | Hoechst 33342 | - | Cell Signaling Technology | 4082S |
| 5 | 18 | CD45RO | AB_528823 | Alexa Fluor 488 | UCHL1 | BioLegend | 304212 |
| 5 | 19 | CD11c | | Alexa Fluor 555 | D3V1E | Cell Signaling Technology | 77882BC |
| 5 | 20 | PD-L1 | AB_2728832 | Alexa Fluor 647 | E1L3N | Cell Signaling Technology | 15005 |
| 6 | 21 | DNA | AB_10626776 | Hoechst 33342 | - | Cell Signaling Technology | 4082S |
| 6 | 22 | CD68 | AB_2798886 | Alexa Fluor 488 | D4B9C | Cell Signaling Technology | 24850 |
| 6 | 23 | FoxP3 | AB_2573608 | eFluor 570 | 236A/E7 | eBioscience/Thermo Fisher | 41-4777-80 |
| 6 | 24 | PD-1 | AB_2728811 | Alexa Fluor 647 | EPR4877(2) | Abcam | ab201825 |
| 7 | 25 | DNA | AB_10626776 | Hoechst 33342 | - | Cell Signaling Technology | 4082S |
| 7 | 26 | CD20 | AB_10734358 | Alexa Fluor 488 | L26 | eBioscience/Thermo Fisher | 53-0202-82 |
| 7 | 27 | Phospho-histone H3.1 (Ser10) | AB_10694639 | Alexa Fluor 555 | D2C8 | Cell Signaling Technology | 3475S |
| 7 | 28 | CD31 | AB_2857973 | Alexa Fluor 647 | EPR3094 | Abcam | ab218582 |
| 8 | 29 | DNA | AB_10626776 | Hoechst 33342 | - | Cell Signaling Technology | 4082S |
| 8 | 30 | E-cadherin | AB_10691457 | Alexa Fluor 488 | 24E10 | Cell Signaling Technology | 3199 |
| 8 | 31 | Vimentin | AB_10859896 | Alexa Fluor 555 | D21H3 | Cell Signaling Technology | 9855 |
| 8 | 32 | Catenin beta-1 | AB_10691326 | Alexa Fluor 647 | L54E2 | Cell Signaling Technology | 4627 |
| 9 | 33 | DNA | AB_10626776 | Hoechst 33342 | - | Cell Signaling Technology | 4082S |
| 9 | 34 | CD163 | AB_2889155 | Alexa Fluor 488 | EPR14643-36 | Abcam | ab218293 |
| 9 | 35 | Histone H3.1 | AB_2799990 | phycoerythrin | D1H2 | Cell Signaling Technology | 82241 |
| 9 | 36 | Phospho-Histone H2AX (Ser139) | AB_2114994 | Alexa Fluor 647 | 2F3 | BioLegend | 613407 |
