## Supplementary Table 4 for "Stitching and registering highly multiplexed whole slide images of tissues and tumors using ASHLAR"

**SUPPLEMENTARY TABLE 4. Description of datasets analyzed.**

| Name | Figures | Tile Dimensions | Grid Dimensions | Pixel Size (µm) | Method | Instrument | Tissue | Source | Location |
|---|---|---|---|---|---|---|---|---|---|
| Colon | 4,5,S1 | 1280 x 1080 | 29 x 21 | 0.65 | t-CyCIF | RareCyte Cytefinder | human colon | Published as part of this work | https://dx.doi.org/10.7303/syn25826362 |
| CRC | 6B | 2048 x 2048 | 15 x 16 | 0.325 | H&E brightfield | GE INCell 6000 | human colorectal adenocarcinoma | Unpublished | |
| Tonsil1 | S2 | 2752 x 2208 | 4 x 4 | 0.227 | CODEX | Zeiss Axio Observer Z1 | human tonsil | Synapse (MCMICRO exemplar data) | https://www.synapse.org/#!Synapse:syn24849819 |
| Spleen | S3 | 1920 x 1440 | 9 x 9 | 0.38 | CODEX | Keyence BZ-X800 | human spleen | HuBMAP (University of Florida TMC / J. P. Aponte) | https://dx.doi.org/10.35079/HBM355.JDLK.244 |
| TMA | S4 | 1280 x 1080 | 36 x 29 | 0.65 | t-CyCIF | RareCyte Cytefinder | human, various | Synapse (EMIT exemplar data) | https://www.synapse.org/#!Synapse:syn22345748/wiki/609239 |
| Tonsil2 | S5 | 1280 x 1080 | 15 x 25 | 0.65 | t-CyCIF | RareCyte Cytefinder | human tonsil | Unpublished | |
| Brain2 | S6 | 2048 x 2048 | 22 x 15 (approximate) | 0.325 | cyclic IF | Zeiss Axio Imager Z2 | rat brain | Figshare (Maric et al.) | https://doi.org/10.6084/m9.figshare.13731585.v1 |
