## Supplementary Figures S1-S5 for "Stitching and registering highly multiplexed whole slide images of tissues and tumors using ASHLAR"

### Figure S1: Registration accuracy comparison on t-CyCIF human colon

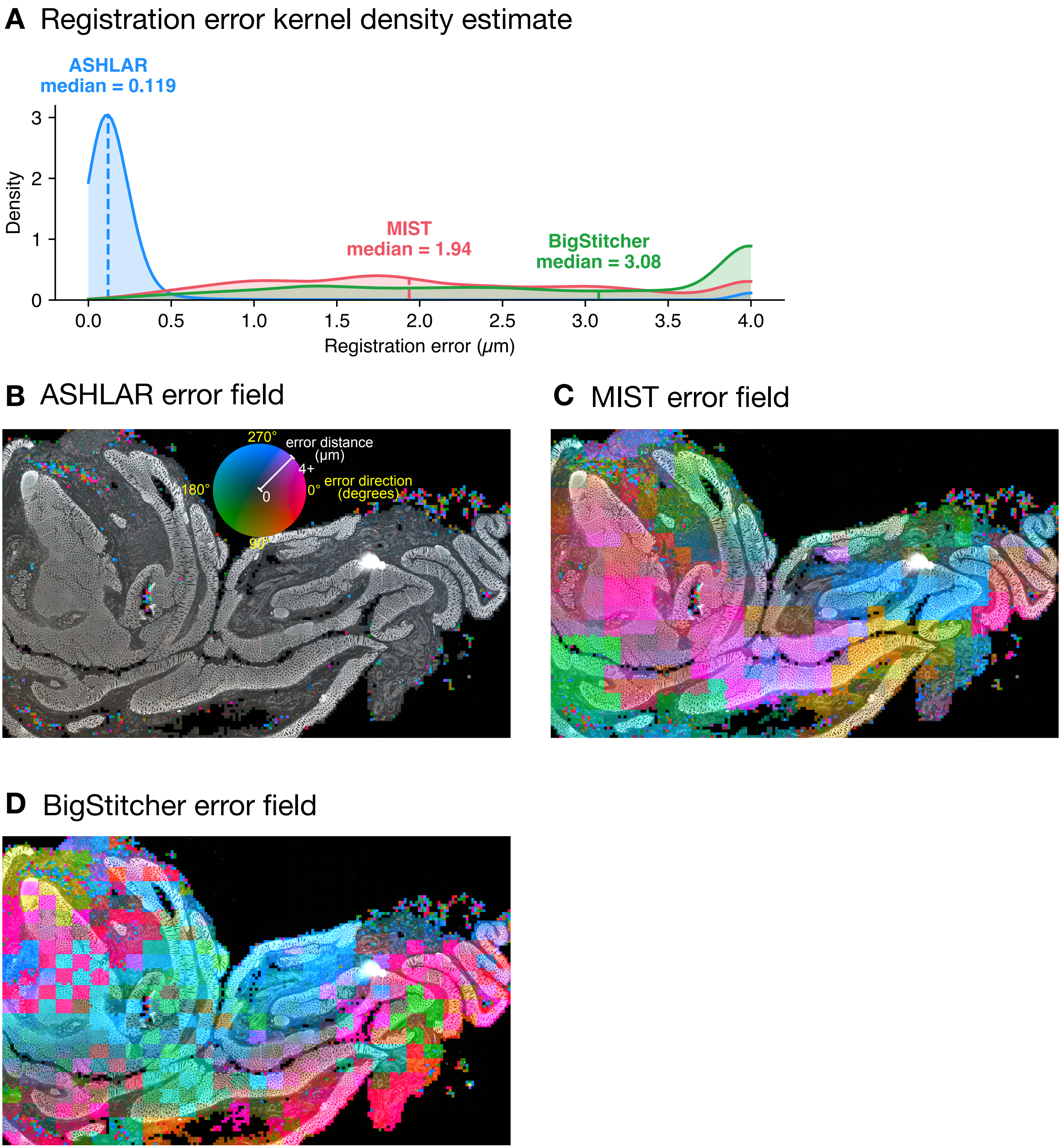


**(A)** Local registration error distance distributions for ASHLAR, MIST, and BigStitcher mosaic images of two t-CyCIF cycles of the human colon section described in the manuscript. Distances at the upper end in this plot as well as in panels B-D were clipped to 4 $\mu$m to highlight the relevant data. **(B-D)** Heatmaps of local registration error for the three tools. Direction (hue) and magnitude (intensity) at 200-pixel resolution are overlaid on the nuclear stain image.

### Figure S2: Registration accuracy comparison on CODEX human tonsil

#### A Registration error kernel density estimate

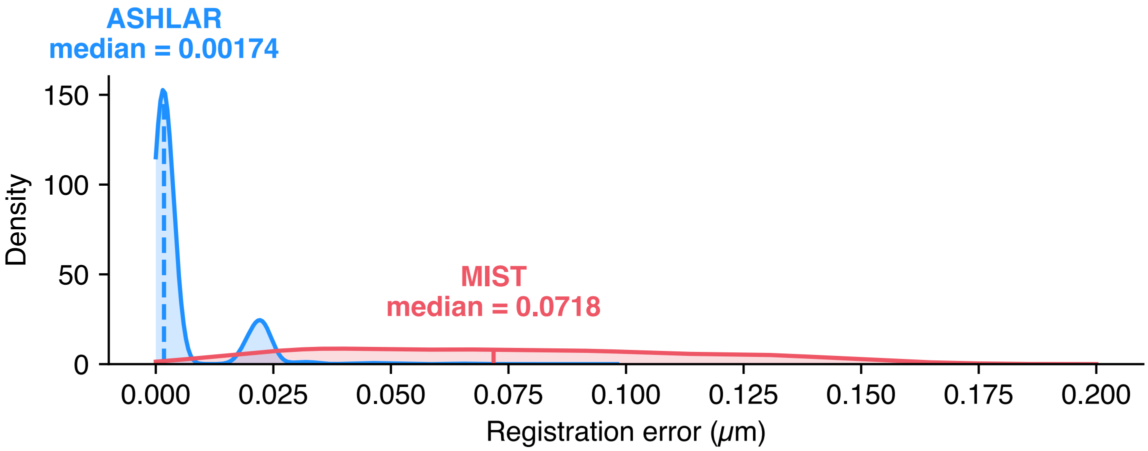


#### B ASHLAR error field

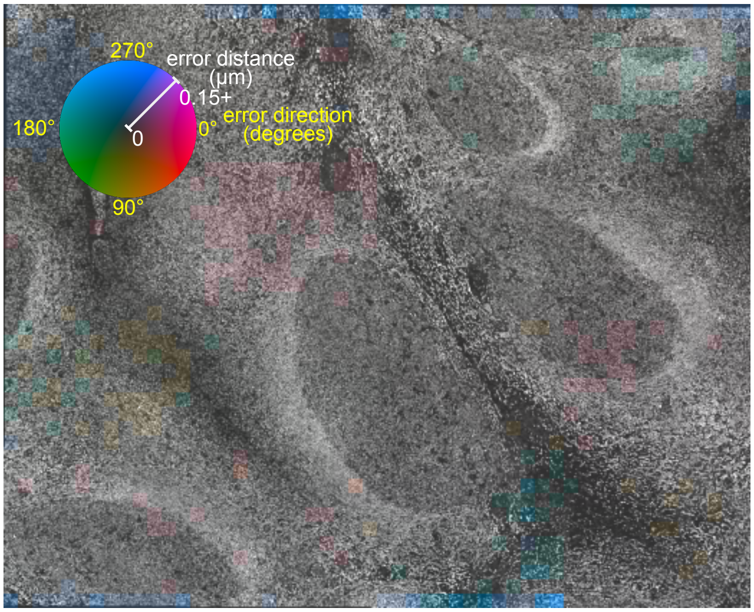


#### C MIST error field

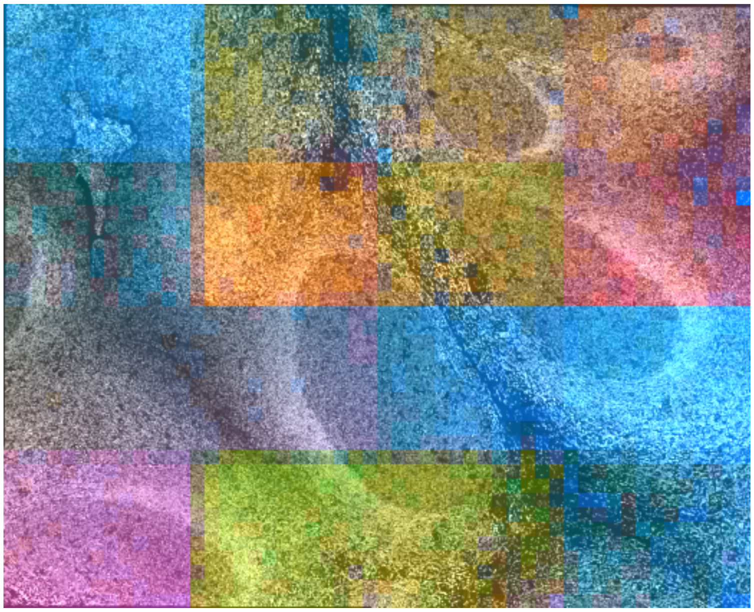

**(A)** Local registration error distance distributions for ASHLAR and MIST mosaic images of two CODEX cycles of a human tonsil section . Distances at the upper end in this plot as well as in panels B and C were clipped to 0.2 $\mu$m to highlight the relevant data. **(B,C)** Heatmaps of local registration error for the two tools. Direction (hue) and magnitude (intensity) at 200-pixel resolution are overlaid on the nuclear stain image.

### Figure S3: Registration accuracy comparison on CODEX human spleen

#### A Registration error kernel density estimate

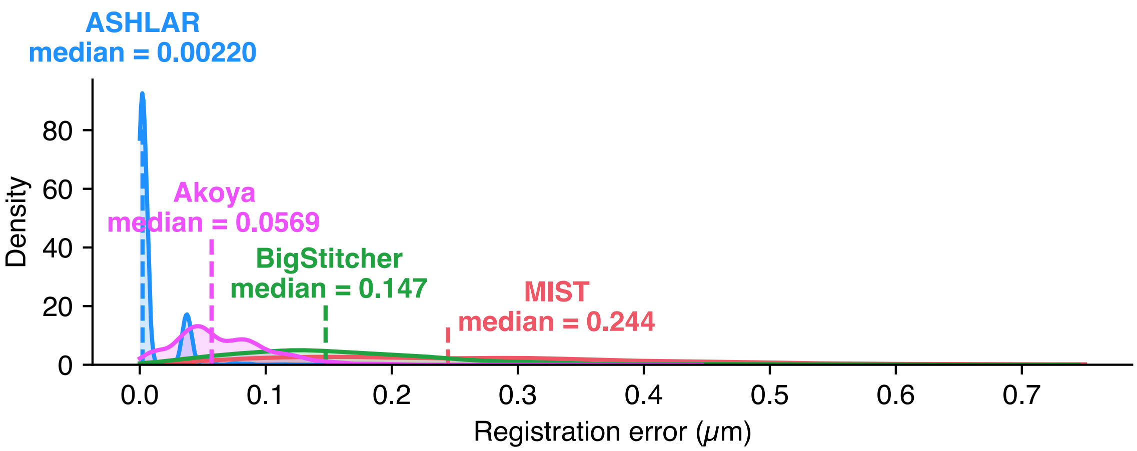


#### B ASHLAR error field

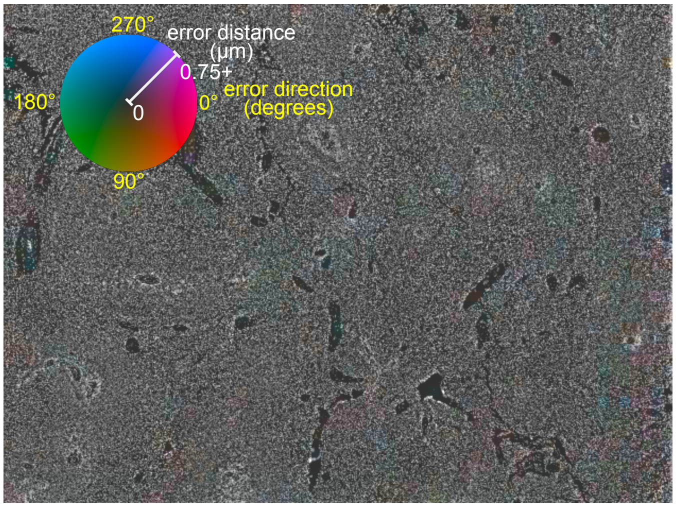


#### C MIST error field

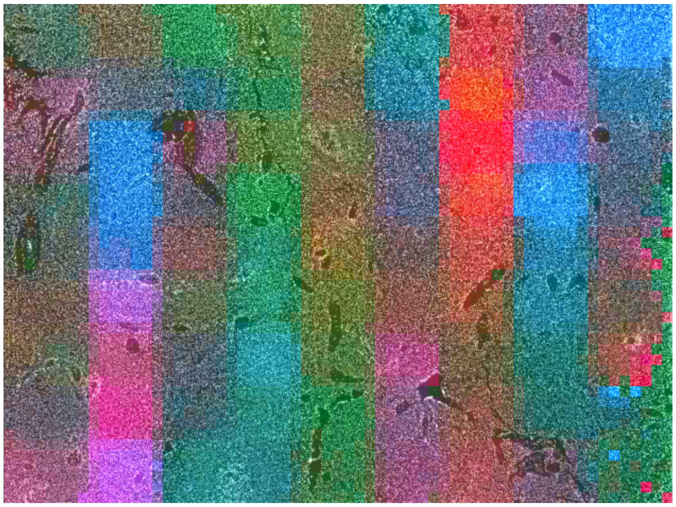

#### D BigStitcher error field

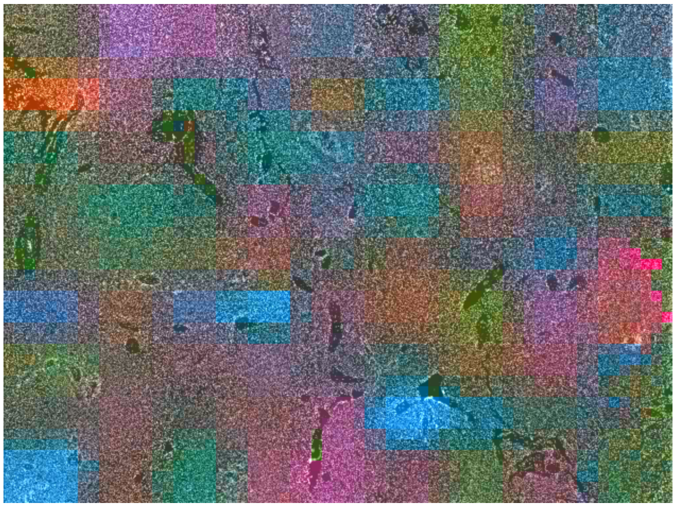

#### E Akoya CODEX pipeline error field

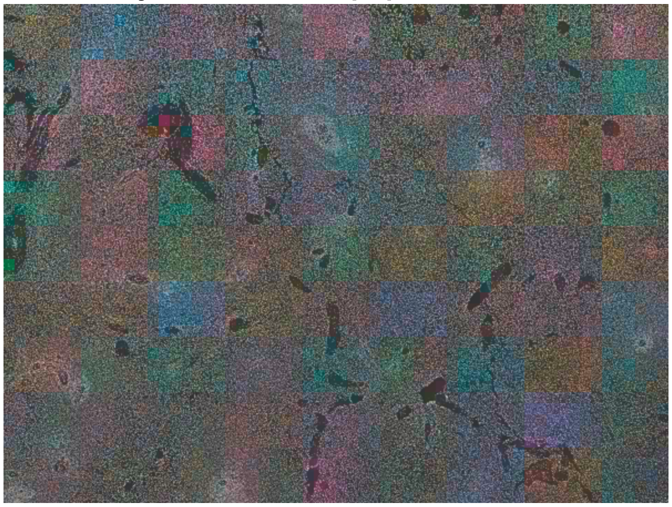

**(A)** Local registration error distance distributions for mosaic images from ASHLAR, MIST, BigStitcher, and the proprietary Akoya CODEX Processor of two CODEX cycles of a human spleen section. Distances at the upper end in this plot as well as in panels B-E were clipped to 0.75 $\mu$m to highlight the relevant data. **(B-E)** Heatmaps of local registration error for the four tools. Direction (hue) and magnitude (intensity) at 200-pixel resolution are overlaid on the nuclear stain image.

### Figure S4: Registration accuracy comparison on CyCIF human TMA

#### **A** Registration error kernel density estimate

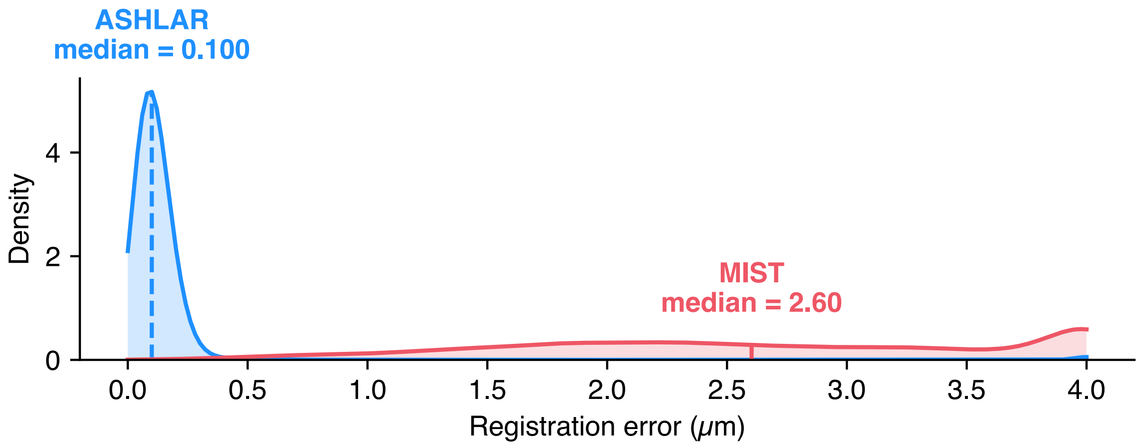


#### **B** ASHLAR error field

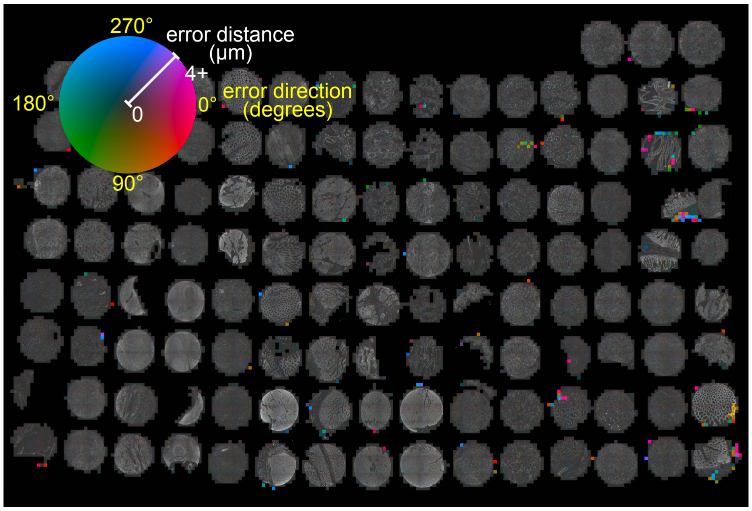


#### **C** MIST error field

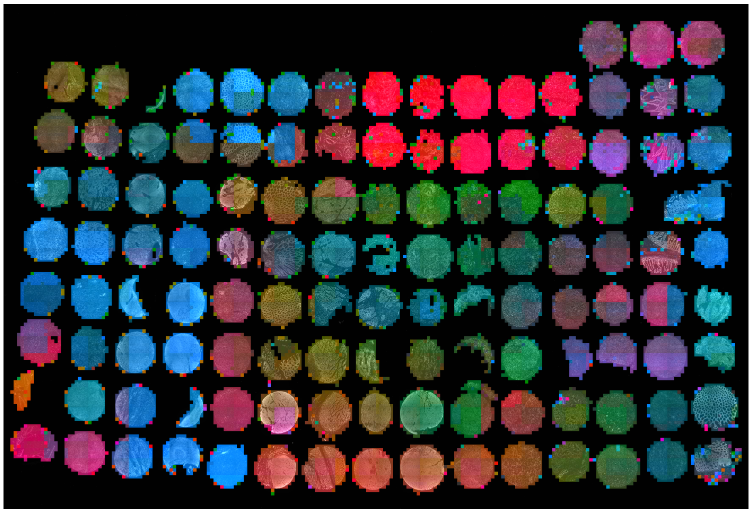

**(A)** Local registration error distance distributions for mosaic images from ASHLAR and MIST of two t-CyCIF cycles of a human tissue microarray (TMA). Distances at the upper end in this plot as well as in panels B and C were clipped to 4 $\mu$m to highlight the relevant data. **(B,C)** Heatmaps of local registration error for the two tools. Direction (hue) and magnitude (intensity) at 200-pixel resolution are overlaid on the nuclear stain image.

### Figure S5: Registration accuracy comparison on CyCIF human tonsil

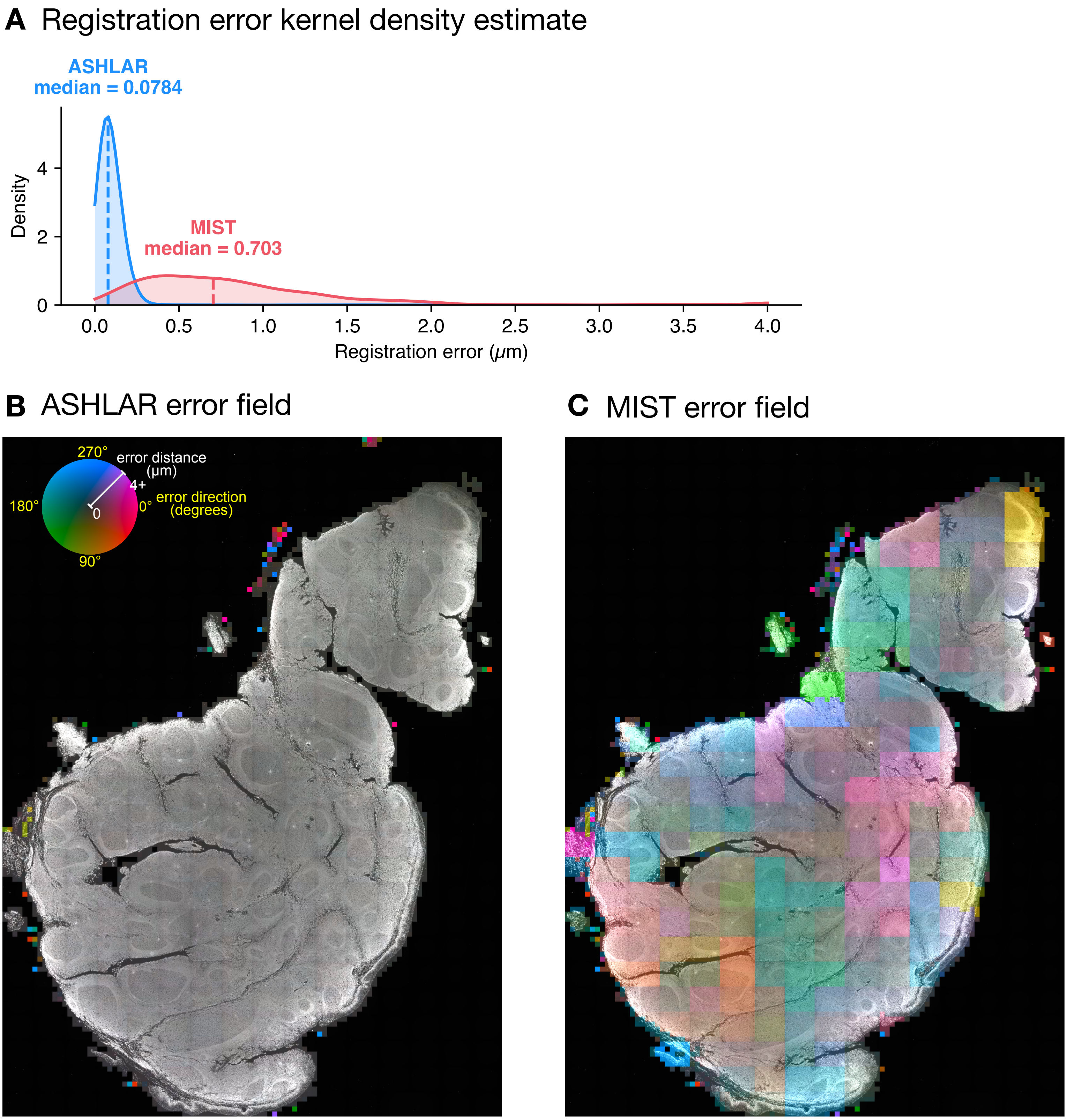


**(A)** Local registration error distance distributions for mosaic images from ASHLAR and MIST of two t-CyCIF cycles of a human tonsil section. Distances at the upper end in this plot as well as in panels B and C were clipped to 4 $\mu$m to highlight the relevant data. **(B,C)** Heatmaps of local registration error for the two tools. Direction (hue) and magnitude (intensity) at 200-pixel resolution are overlaid on the nuclear stain image.
