## Supplementary Figure S6 for "Stitching and registering highly multiplexed whole slide images of tissues and tumors using ASHLAR"

**Figure S6: Registration accuracy of Zeiss Zen software on cyclic IF rat brain**

**A** Registration error kernel density estimate

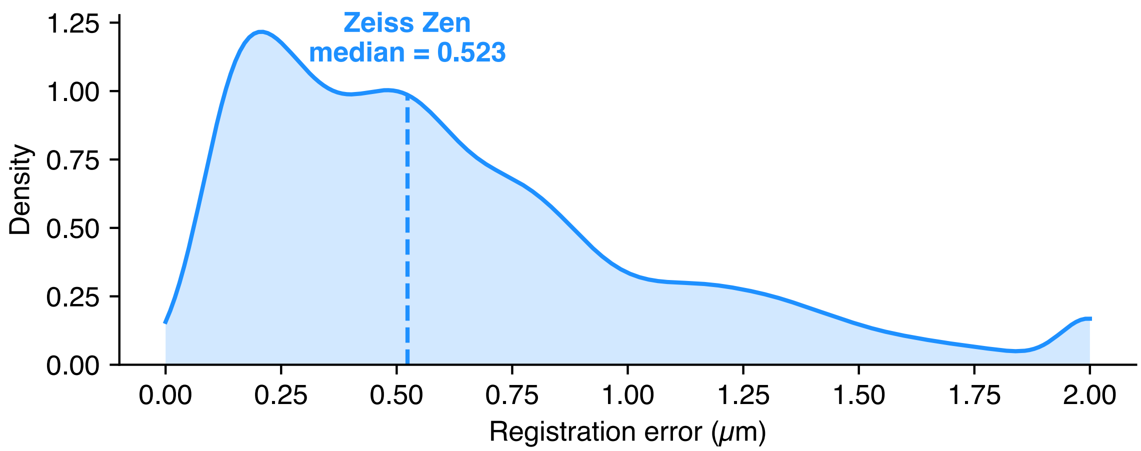


**B** Zeiss Zen error field

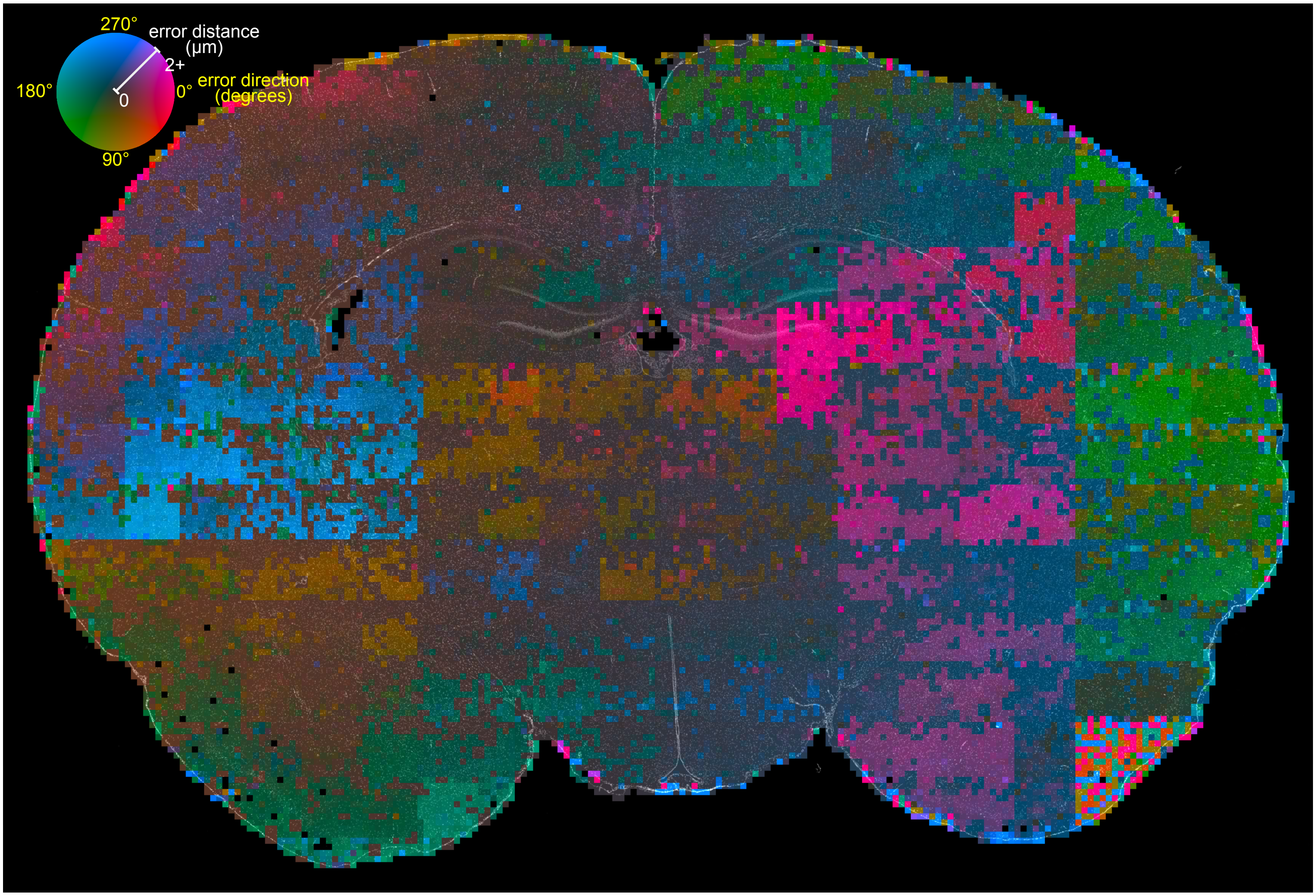


**(A)** Local registration error distance distributions for mosaic images from Zeiss Zen software of two cyclic IF cycles of a rat brain section. Distances at the upper end in this plot as well as in panel B were clipped to 2 $\mu$m to highlight the relevant data. **(B)** Heatmap of local registration error. Direction (hue) and magnitude (intensity) at 200-pixel resolution are overlaid on the nuclear stain image.
