## Supplementary Figure S7 for "Stitching and registering highly multiplexed whole slide images of tissues and tumors using ASHLAR"

### Figure S7: ASHLAR runtime vs. dataset size

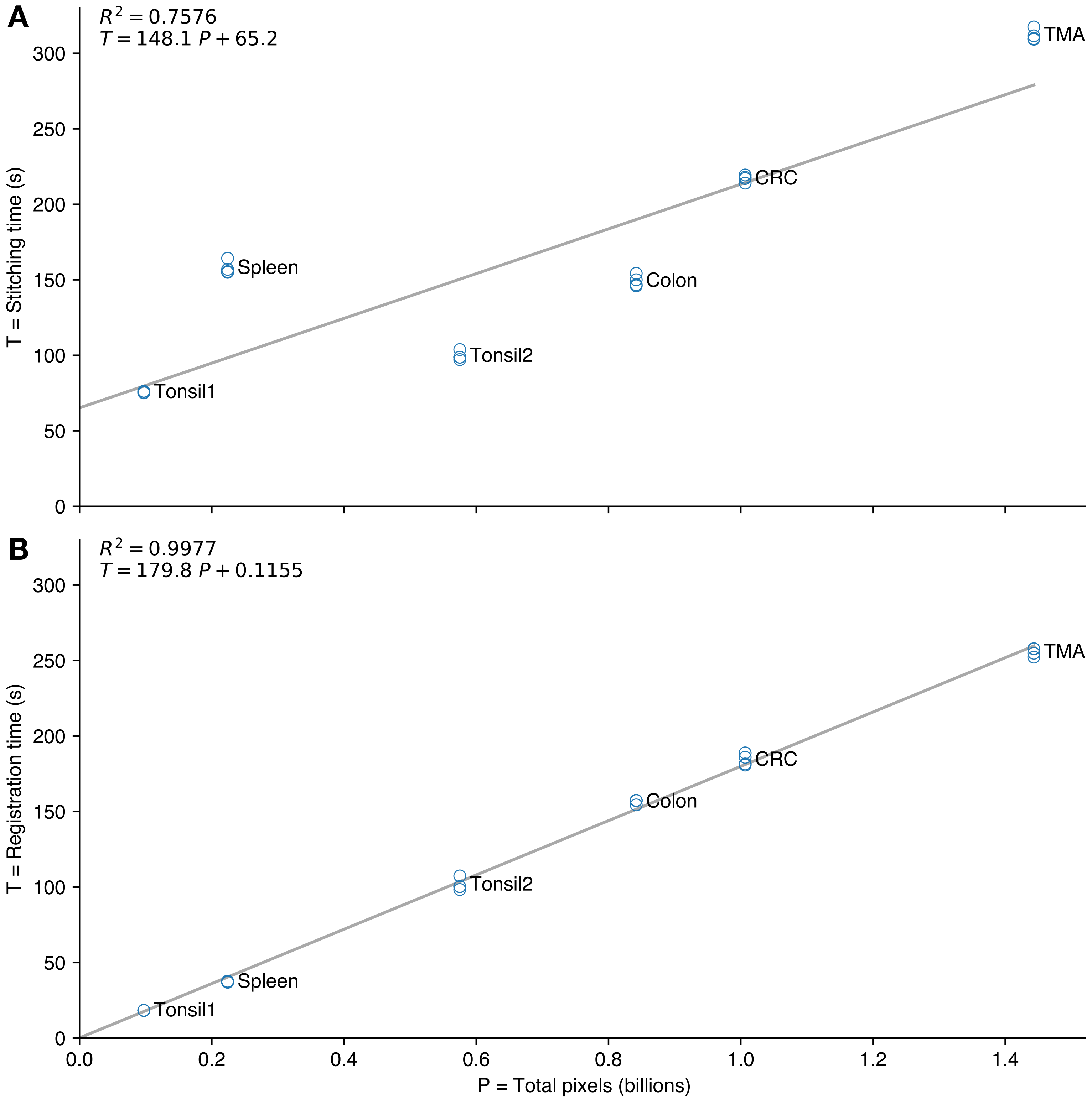


ASHLAR runtime for the stitching phase **(A)** and registration phase **(B)** vs. dataset size in total pixels per imaging cycle. Data points are shown for four runs of each of six datasets along with a linear fit. Datasets are summarized in **Supplementary Table 4**.
